# The functional impact of rare variation across the regulatory cascade

**DOI:** 10.1101/2022.09.07.507008

**Authors:** Taibo Li, Nicole Ferraro, Benjamin J. Strober, Francois Aguet, Silva Kasela, Marios Arvanitis, Bohan Ni, Laurens van de Wiel, Elliot Hershberg, Kristin Ardlie, Dan E. Arking, Rebecca L. Beer, Jennifer Brody, Thomas W Blackwell, Clary Clish, Stacey Gabriel, Robert Gerszten, Xiuqing Guo, Namrata Gupta, W. Craig Johnson, Tuuli Lappalainen, Henry J. Lin, Yongmei Liu, Deborah A. Nickerson, George Papanicolaou, Jonathan K. Pritchard, Pankaj Qasba, Ali Shojaie, Josh Smith, Nona Sotoodehnia, Kent D. Taylor, Russell P. Tracy, David Van Den Berg, Matthew Wheeler, Stephen S. Rich, Jerome I. Rotter, Alexis Battle, Stephen B. Montgomery

**Author notes:** co-first authors (Taibo Li, Nicole Ferraro). co-senior authors (Alexis Battle, Stephen B. Montgomery).

## Abstract

Each human genome has tens of thousands of rare genetic variants; however, identifying impactful rare variants remains a major challenge. We demonstrate how use of personal multi-omics can enable identification of impactful rare variants by using the Multi-Ethnic Study of Atherosclerosis (MESA) which included several hundred individuals with whole genome sequencing, transcriptomes, methylomes, and proteomes collected across two time points, ten years apart. We evaluated each multi-omic phenotype’s ability to separately and jointly inform functional rare variation. By combining expression and protein data, we observed rare stop variants 62x and rare frameshift variants 216x as frequently as controls, compared to 13x to 27x for expression or protein effects alone. We developed a Bayesian hierarchical model to prioritize specific rare variants underlying multi-omic signals across the regulatory cascade. With this approach, we identified rare variants that exhibited large effect sizes on multiple complex traits including height, schizophrenia, and Alzheimer’s disease.

## Introduction

There are thousands of rare (minor allele frequency; MAF < 1%) genetic variants in every human genome but determining which, if any, exert a significant phenotypic effect remains challenging. Prior work has demonstrated the use of transcriptome data in prioritizing rare variants with both large molecular and phenotypic effects^1,2^. However, rare variants have the potential to influence additional regulatory mechanisms beyond transcription, such as DNA methylation and protein expression, and integrating corresponding functional genomics data can allow for more comprehensive detection of impactful rare variants and understanding of their roles in the regulation of gene function.

The ability of transcriptome data to enhance prioritization of rare variants with effects on diseases and traits^3^ is presumably due to those effects propagating through the regulatory cascade to protein levels and cellular functions. Prior work has shown that common variants associated with changes in gene expression can have effects on ribosome and protein levels, though those effects are significantly reduced at the protein level^4,5^. We and others have also shown that common variants can be associated with changes in protein abundance, yet not show any impact at the mRNA level, indicating the effects of post-translational regulation, in addition to the substantial effects of post-transcriptional and protein degradation regulation^4–6^. In particular, the plasma proteome contains proteins generated from many different cell types, leading to its regular use as a source for biomarker discovery^6,7^; therefore, understanding how rare genetic variation impacts protein abundance in samples such as plasma may help identify impactful rare variants from tissues that are more challenging to transcriptome-sequence^8^.

In this study, we expand the assessment of impactful rare variation to integrate molecular signatures across the regulatory cascade. We analyzed measurements of DNA methylation from whole blood, RNA-sequencing from peripheral blood mononuclear cells (PBMCs), and plasma proteome abundance from a multi-ethnic cohort of ∼900 individuals with data from two time points, ten years apart, and assessed the ability of each measurement to prioritize nearby rare variation. Notably, we observed that the longitudinal design of these data provided robust outlier measurements per individual per data modality. We subsequently integrated these diverse functional signals into a predictive model to assign probabilities to individual rare variants leading to functional effects at various levels of the regulatory cascade. Finally, we demonstrated the utility of these predicted functional probabilities in prioritizing variants with large effects on downstream traits and diseases.

## Results

### Consistency of outlier measurements across time

Assessing the correlation of multi-omic measurements across between ten-year timepoints of collection, plasma proteome measurements exhibited the highest correlation (R = 0.64), followed by expression (R = 0.27), gene-level methylation (R = 0.20), and gene-level splicing (R = 0.07) (Figure S1). We then assessed replication across time for the subset of measurements at the extremes of the distribution (“outliers”) for each gene-level outlier type. We refer to those instances for gene expression as “eOutliers”, methylation as “mOutliers”, splicing as “sOutliers”, and protein as “pOutliers”. After identifying outliers in exam 1, based on an individual’s deviation from the mean for a given gene (Z-score), we assessed the proportion that also had measurements at least two standard deviations from the mean in exam 5. Across thresholds, we observed the highest replication for pOutliers (range = [0.34 - 0.89]), followed by mOutliers (range = [0.18 - 0.82]), eOutliers (range = [0.12 - 0.85]), and then sOutliers (range = [0.07,0.22]). When focusing on the subset of eOutliers with negative Z-scores and thus very low expression, we saw the replication rate across time increased with threshold stringency (Figure 1A), eventually surpassing all other replication rates when the measurements were over ∼6 standard deviations below the mean (Z < -6), at which point 79% of exam 1 eOutliers were also seen in exam 5. This is in line with prior work demonstrating that under-expression outliers are more often associated with rare variants, and thus likely more often genetically-driven than over-expression outliers^1,2^. In order to focus on robust and more likely genetically-driven outlier events, we took advantage of the longitudinal study design and required an outlier effect to be seen in both time points in subsequent analyses (“joint outliers”). For joint outliers, we observed an average of 12.5 eOutliers, 1.2 gene-level mOutliers (472 CpG-level mOutliers), 4.8 sOutliers (9.9 sOutlier clusters), and 8.3 pOutliers per individual (Figure 1B).

**Figure 1.**
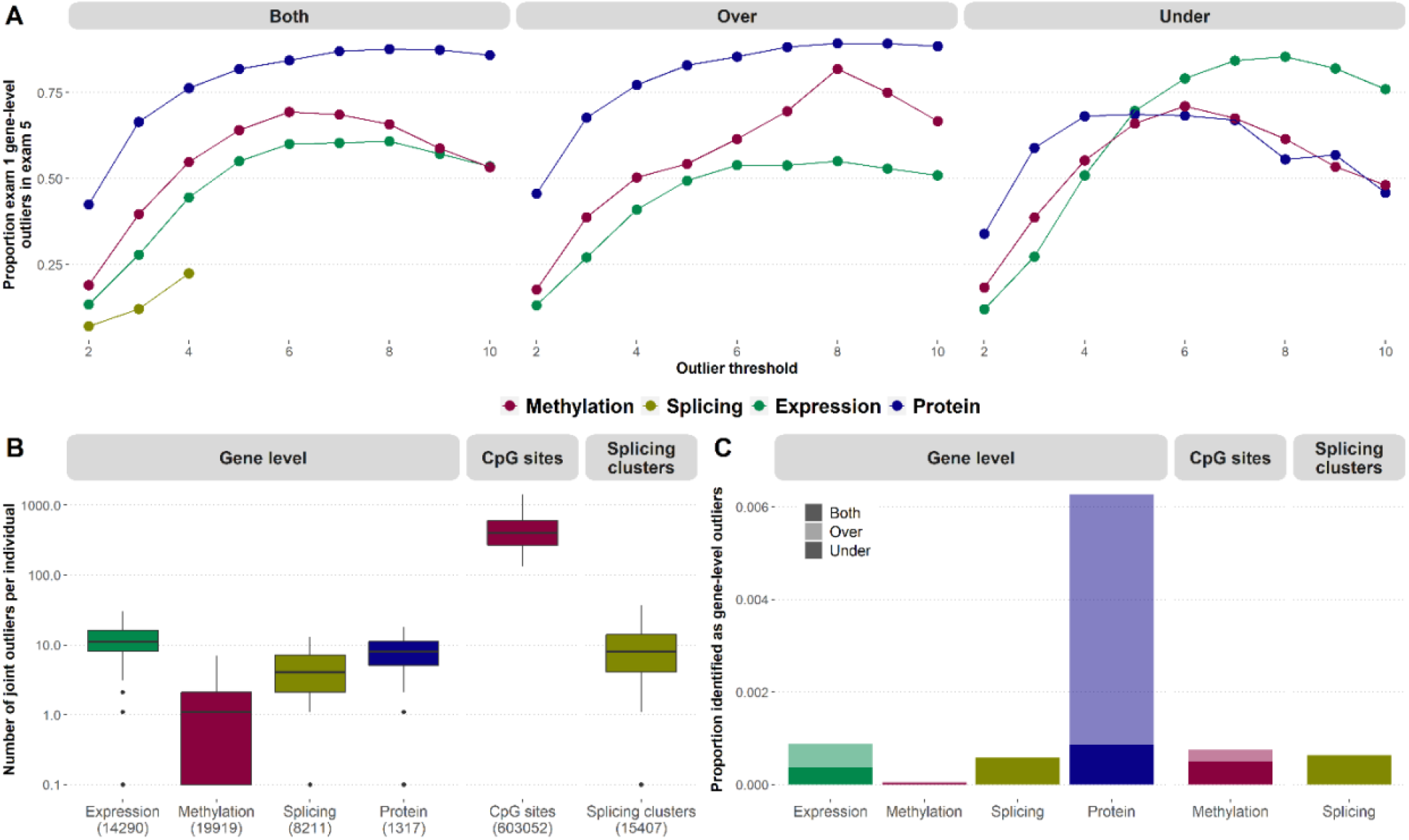
Outlier calls across exams. **(A)** The proportion of gene-level outliers identified in exam 1 (y-axis) at varying thresholds (x-axis) that replicate in exam 5 at a threshold of |Z| > 2. sOutliers (gold) do not have direction and so are shown only for the combined set of outlier calls (left), while eOutliers (green), mOutliers (red), and pOutliers (blue) are also shown split by direction, with outliers with positive Z-scores in the center (“Over”) and negative Z-scores on the right (“Under”). **(B)** The number of outliers identified per individual where the outlier effect is seen in both exams, using a threshold of |Z| > 3 for all gene-level outliers (left), as well as the number of CpG-level mOutliers (center) and sOutlier clusters (right). **(C)** The proportion of all gene-individual pairs considered that show outlier signal in both exams, using a threshold of |Z| > 3, split by the direction of the effect.

We assessed the relative proportion of joint outlier events for each omics data type. Restricting to individuals with data in both time points, we assessed 14,290 genes across 547 individuals for eOutliers, 19,919 genes across 785 individuals for mOutliers, 8,211 genes across 564 individuals for sOutliers, and 1,317 proteins across 876 individuals for pOutliers. Looking at the proportion of tests that resulted in joint outliers at a threshold of |Z| > 3, we found the highest proportion for pOutliers, followed by eOutliers (Figure 1C). Overall, the set of pOutliers contained many more high abundance outliers compared to low abundance outliers, while the proportions in either direction were more comparable for eOutliers and mOutliers. This could reflect the dynamic range of the protein measurements, as previous work has found the range of protein abundances detectable to be higher than that of mRNA transcripts^9^, and there is no strict upper bound for high abundance outliers, while detected protein abundances can only decrease to 0. We further found that the number of high and low abundance pOutliers discovered varies by protein type (Figure S2A) or inferred tissue of origin (Figure S2B) and observed that classes of proteins with higher base expression tended to have more low abundance pOutlier individuals than the set of all proteins and vice versa (Figure S2C-D).

### Outlier sharing across the regulatory cascade

While each data type was measured in different biospecimens, with DNA methylation from whole blood, expression data from peripheral blood mononuclear cells (PBMCs), and protein measurements from plasma, we assessed the sharing of outlier signals across each omics data type, as rare variant effects can manifest across multiple tissues^1,2^. For the set of joint outliers identified in each data type at a threshold of |Z| > 3, we assessed the mean Z-scores across exams in all other gene-level data types. For under-eOutlier individuals, there were significant shifts in corresponding methylation (p = 1.5e-15, one-sided Wilcoxon rank sum test), splicing (p < 2.2e-16), and protein (p = 5.1e-14) Z-scores. For over-eOutlier individuals, there were significant shifts in methylation (*p* < 2.2e-16) and splicing (*p* = 2.5e-5) Z-scores for over-eOutliers (Figure 2A). For gene-level mOutlier and sOutlier individuals, there was a significant increase in the corresponding expression Z-scores (*p* = 2.8e-13 and *p* < 2.2e-16, respectively). For low abundance pOutlier individuals, there was a corresponding significant shift in expression values (*p* = 1.1e-11), though this is not the case for high abundance pOutlier individuals.

**Figure 2.**
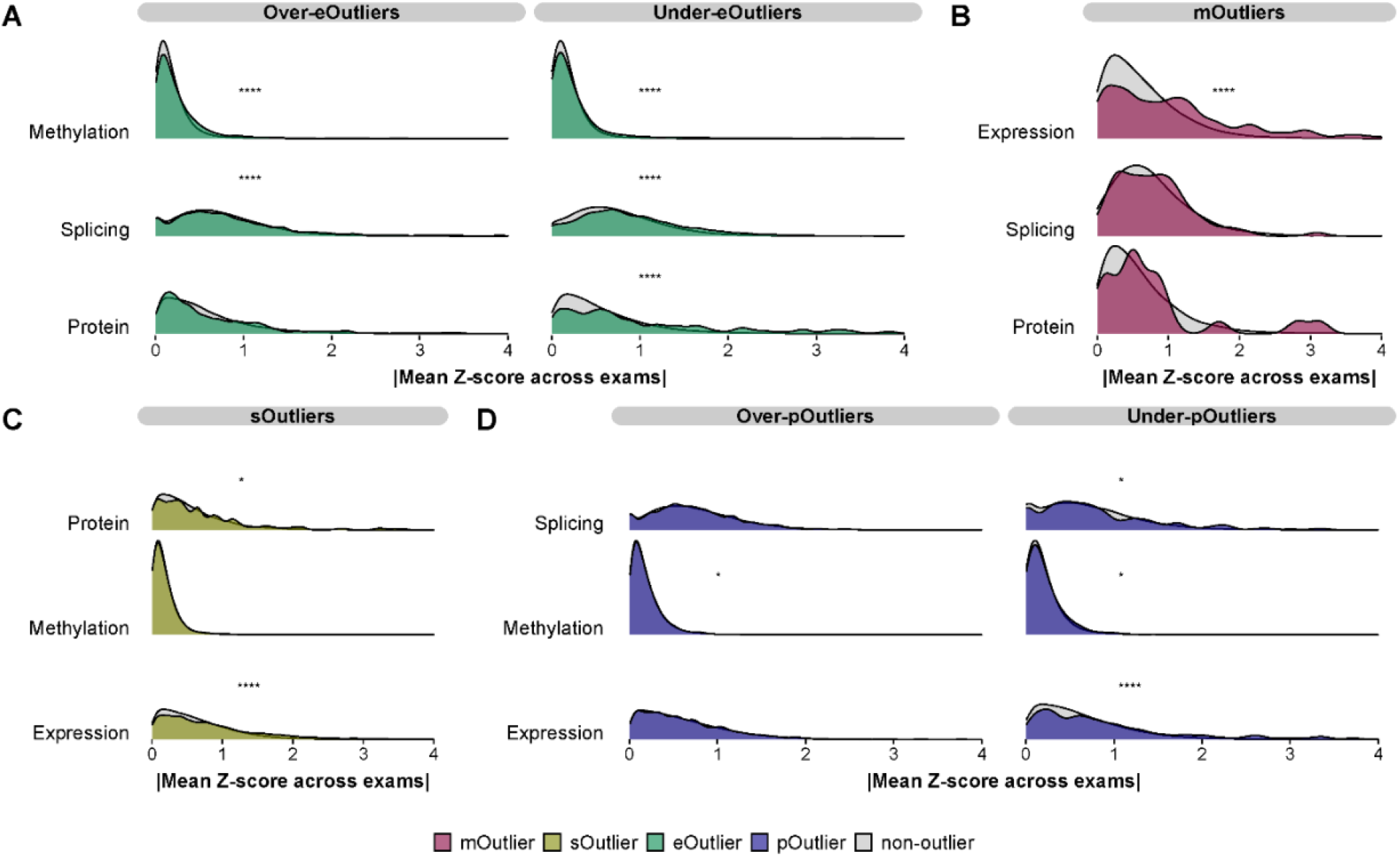
Distribution of Z-scores for outliers in other data types. **(A)** The distribution of gene-level methylation, gene-level splicing, and protein Z-scores for eOutlier individuals (green) and non-outliers (grey) for corresponding genes, split by the direction of the expression effect. **(B)** The distribution of expression, gene-level splicing, and protein Z-scores for mOutlier individuals (red) and non-outliers (grey) for corresponding genes. **(C)** The distribution of expression, gene-level methylation, and protein Z-scores for sOutlier individuals (gold) and non-outliers (grey) for corresponding genes. **(D)** The distribution of expression, gene-level methylation, and gene-level splicing Z-scores for pOutlier individuals (blue) and non-outliers (grey) for corresponding genes, split by the direction of the expression effect. **** p < 0.0001, * p < 0.05, one-sided Wilcoxon rank-sum test on absolute value of mean Z-score across both exams between outlier and non-outlier individuals.

For eOutliers, the highest degree of sharing was seen at the protein level. Of 485 eOutliers (|Z| > 3 in both exams) identified in genes and individuals that also had protein measurements, we found that 12% of those (N = 58) were shared at the protein level, with 29.3% (N = 17) of those being high abundance pOutliers. For all other gene-level outlier types (mOutliers, sOutliers, and pOutliers), the highest degree of sharing was seen at the expression level, with 18.2%, 8.9%, and 3.7% of mOutliers, sOutliers, and pOutliers, respectively (Figure S3A). Considering only under-eOutliers, 20.8% of those showed outlier protein levels and for low abundance pOutliers, 15.1% also had outlier expression levels (Figure S3B). Overall, eOutliers had the strongest shift in values for other functional measurements and best captured outlier signals across all other data types, particularly when the outlier effect led to very low expression, indicating that the transcriptome best captured effects that propagated throughout the regulatory cascade, though any one individual measurement will also miss some potentially abnormal function.

### Rare variants contribute to outlier effects across multi-omics data types

We expect rare variants to contribute substantially to the observed outlier effects, as has been thoroughly demonstrated for transcriptome outliers^2^ and investigated for methylation^10,11^ and protein levels^12^. As we have multiple -omics measurements for the same individuals across this cohort, and observed that a proportion of outlier effects are shared between molecular phenotypes, we sought to assess the degree to which rare variation contributed to each gene-level outlier signal and the benefit of collecting multiple-omics measurements for rare variant interpretation.

Considering each of the four gene-level outlier types individually, we observed the strongest enrichment for mOutliers, which carried rare variants in the outlier gene body or within 10kb between 1.11x to 1.55x as frequently as non-outliers, depending on threshold stringency. This was followed by eOutliers (relative risk = [1.10 - 1.29]), sOutliers (relative risk = [1.02 - 1.26]) and pOutliers (relative risk = [1.03 - 1.06]), considering rare variants within the same 10kb-window. pOutliers had the smallest enrichment despite having highest replication across exams, though a recent study looking at common variants impacting plasma proteome (pQTLs) reported ∼60% of proteins had only trans-pQTLs (>500kb from target), suggesting that impactful rare variants may be more often located in trans. For joint CpG-level mOutliers, they were strongly enriched for carrying nearby rare variants across windows that ranged from 100bp (relative risk = 52.9, *p* < 2.2e-16) to 1kb (relative risk = 5.84, *p* < 2.2e-16) around the site. These enrichments remained significant but were largely driven by instances where rare variants overlapped the CpG site itself (Figure S4A-B). As a further signature of a rare variant effects, CpG-level mOutliers also showed a significant increase in allele-specific expression in a 1kb window around outlier sites (Figure S4C).

Previous studies have observed stronger rare variant enrichments for under-eOutliers compared to over-eOutliers^2^. For expression, we observed similar patterns (relative risk = [1.12 - 1.37] for under-eOutliers and [1.10 - 1.24] for over-eOutliers). This observation also held for other omics data types included in our study. When splitting by the direction of the effect for methylation, hypo-methylated mOutliers (relative risk = [1.13 - 1.73]) showed stronger enrichments than hyper-methylated mOutliers (relative risk = [1.13 - 1.29]). Likewise, for pOutliers we observed higher enrichment for under-outliers (relative risk = [1.19 - 1.34]) compared to over-outliers (relative risk = [0.99 - 1.02]). Notably, high abundance pOutliers were not significantly enriched for nearby rare variation at any threshold above |Z| > 2 (relative risk = 1.02, *p* = 1.54e-3) (Figure 3A). This lack of enrichment was not entirely due to the restricted set of genes assayed for protein abundance as restricting the set of eOutliers to the genes also assayed at the protein level, they were still significantly enriched for nearby rare variants in the over-expression direction at thresholds of |Z| > 2 (relative risk = 1.10, *p* = 1.97e-8) and |Z| > 3 (relative risk = 1.11, *p* = 0.013; Figure S5).

**Figure 3.**
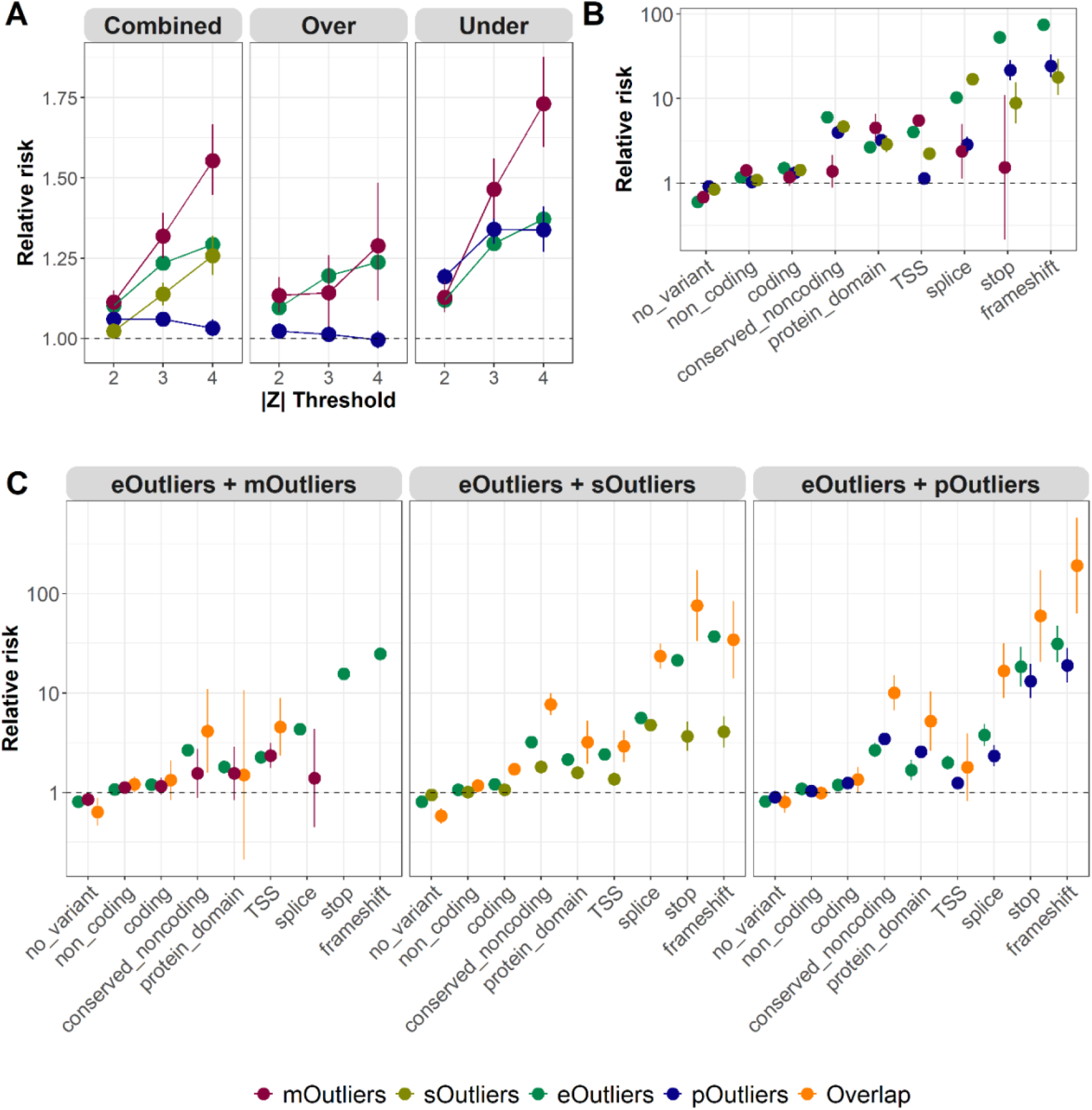
Enrichment of rare variants nearby gene-level outliers. **(A)** The relative risk of nearby rare variants for eOutliers (green), gene-level mOutliers (red), gene-level sOutliers (gold), and pOutliers (blue) across varying Z-score thresholds (x-axis). Enrichments are split by the direction of the effect for eOutliers, mOutliers, and pOutliers. Non-outliers are defined as all individuals with |Z| < 1 in both exams for the same set of genes. **(B)** The relative risk of nearby rare variants with a given annotation (x-axis) for eOutliers (green), gene-level mOutliers (red), gene-level sOutliers (gold), and pOutliers (blue) at a threshold of |Z| > 3 in both exams. Non-outliers are defined as all individuals with |Z| < 1 in both exams for the same set of genes. If no outlier individual carried a nearby rare variant in a given category, that data type is not shown. **(C)** The relative risk of nearby rare variants with a given annotation (x-axis) for combinations of eOutliers and the other three data types (orange) as compared to single data type outliers, matched for the same considered genes and individuals, considering a reduced threshold of |Z| > 2 in both exams, and in both data types for the overlapping set. Non-outliers are defined as all individuals with |Z| < 1 in both exams for the same set of genes. If no outlier individual carried a nearby rare variant in a given category, that data type is not shown.

We next evaluated if different categories of rare variants contributed to observed enrichments in each omics data type. Considering different predicted effects across variants, eOutliers were most strongly enriched for nearby rare stop and frameshift variants, as expected based on previous work^2^, while sOutliers were most strongly enriched for nearby rare splice variants, with strong enrichments also for rare stop and frameshift variants. For gene-level mOutliers, most variant categories were not significantly enriched, but rare variants nearby the associated gene’s transcription start site (TSS) were seen 5.49x (p < 2.2e-16) as frequently in mOutlier individuals as compared to controls (Figure 3B). While the combined set of pOutliers were largely not enriched for nearby rare variants overall, there was strong enrichments for nearby rare stop (relative risk = 21.7, *p* < 2.2e-16) and frameshift (relative risk = 24.4, *p* < 2.2e-16) variants, though this was predominantly driven by under-expression pOutliers (Figure S6).

### Multi-omics outliers increase discovery of rare variant effects

As expression outliers best captured outlier signals in other data types (Figure 2; Figure S3), we assessed the gain in rare variant enrichments when considering eOutliers in conjunction with outliers for each of the other data types. While for many types of variants, eOutliers alone tagged functional rare variants at a similar frequency to the set of multimodal outliers, there was a subset of variant types for which multi-omics data improved functional rare variant identification (Figure 3C). Most notably, the set of gene-individual pairs that showed outlier signal at both the expression and protein level are more strongly enriched for nearby rare conserved non-coding, protein-domain, splice, stop, and frameshift variants as compared to the set of eOutliers or pOutliers identified alone in the same set of genes and individuals. When considering both expression and methylation signal, there was an improvement in enrichment for nearby rare TSS variants over either data type alone, and for overlapping expression and splicing signal, the enrichment of nearby rare conserved non-coding variants and rare splice and stop variants were all increased (Figure 3C), indicating that for specific variant effects, assessing multiple molecular signals can improve identification of functional rare variants.

In practice, it may be difficult to collect both multiple ‘ omics measurements from an individual as well as data across multiple time points. While we are limited by the relatively smaller number of proteins assayed as compared to gene expression measurements, we assessed the relative gain in enrichments considering both expression and protein outliers identified from only a single time point as compared to outlier effects seen in each specific omics data type measured at two time points. While the set of overlapping eOutliers and pOutliers at a single time point is small (n = 72 at a threshold of |Z| > 3), we do see increased enrichment of nearby rare variation (relative risk = 1.37, *p* = 1.71e-4) over either joint eOutliers (relative risk = 1.23, *p* < 2.2e-16) or joint pOutliers (relative risk = 1.06, *p* = 1.72e-9) at that same threshold or higher (Figure S7). This indicates that multi-omics measurements are providing enhanced ability to detect rare variant-driven outliers compared to repeated measures of a specific omics data type.

### Replication of GTEx outlier-associated rare variants

Our previous work identified rare variants associated with multi-tissue transcriptome outliers in the Genotype Tissue Expression project (GTEx)^2^, which consisted primarily of individuals of European ancestry. Here, we observed significant correlation between individual outlier burden and genotype principal components (PCs), which decreased at increasing outlier thresholds (Figure S8). Notably, we saw little difference in rare variant enrichment estimates after matching each outlier individual to a control individual by ancestry, as measured by genotype PCs (Figure S9), and thus did not observe evidence of differences in genetic ancestry driving the observed enrichment of rare variants nearby any outlier type. Next, we evaluated the proportion of those GTEx variants that are carried by any individual in MESA and exhibit consistent effects on gene expression and splicing. For eOutliers, we identified 1348 multi-tissue eOutlier-associated variants in GTEx that were present in any MESA individual and occurred at <1% frequency across MESA, which totaled 5604 total variant-gene-individual instances (Figure S10A). Of these, 888 also showed outlier expression in MESA, at a reduced threshold of |Z| > 2 in both exams (q<0.01; Figure S10B). We found that rare stop variants are most predictive of replicating expression effects, followed by rare splice variants (Figure S10C). For sOutliers, we identified 1113 multi-tissue sOutlier-associated variants in GTEx that were present in any MESA individual, which totaled 5858 total variant-gene-individual instances (Figure S11A).

Of these, 891 also showed outlier splicing in MESA, at a reduced threshold of |Z| > 2 in both exams (q<0.01; Figure S11B). We observed that rare splice variants are most predictive of replicating outlier splicing effects (Figure S11C).

### Development and evaluation of multi-omic rare variant prediction model

To leverage the full spectrum of data available in MESA to prioritize functional rare variants, we extended our Bayesian hierarchical variant effect prediction model, Watershed^2^. Briefly, Watershed integrates genomic annotations such as conservation scores and variant annotations with observed outlier signals from functional data in a latent variable model originally developed for transcriptomic outliers. Here, we extended Watershed to include mRNA expression, methylation, splicing, and protein expression (Figure S12A). When evaluated against pairs of individuals with the same rare variant nearby the same gene (N2 pair), the multi-omic Watershed model outperformed logistic regression models based on genomic annotations alone (GAM) in predicting the regulatory status of one individual in the pair based on the genomic annotations and observed outlier status in the other, achieving an area under the precision-recall curve between 0.07 and 0.11 across the four omics data types (Figure 4A), compared to 0.02 to 0.06 for GAM.

**Figure 4.**
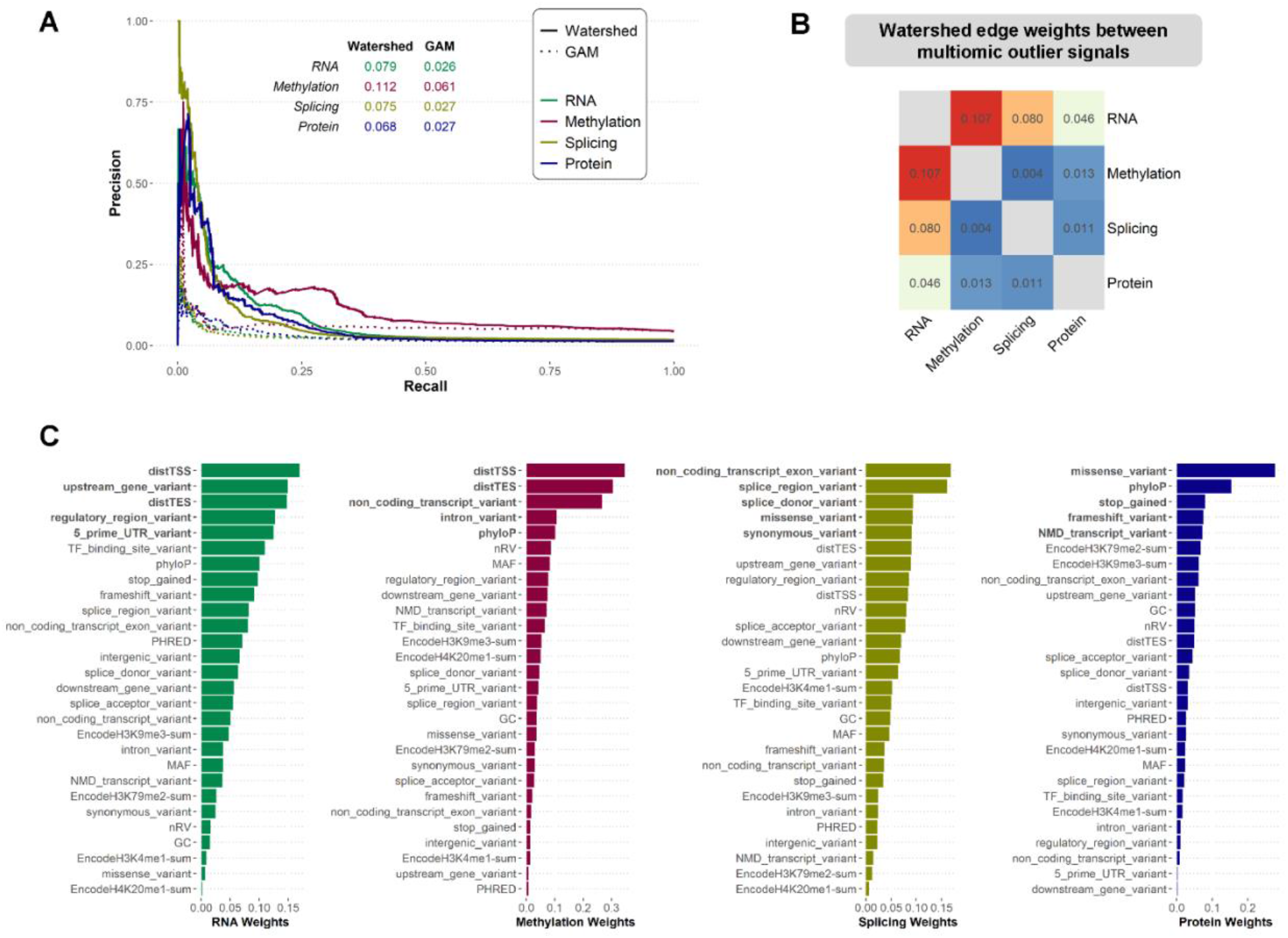
Evaluation of multi-omic Watershed model. **(A)** Precision-recall curves of Watershed models (solid lines) and genomic annotation models (GAM, dotted lines) for mRNA expression (green), methylation (red), splicing (gold), and protein expression (blue) evaluated against (gene, individual) pairs with the same set of rare variants nearby. GAM outliers are defined by a p-value threshold of 0.05. **(B)** Symmetric matrix summarizing weights of edges connecting the latent regulatory variables (Z) in the multi-omic Watershed model. **(C)** Edge weights connecting top genomic annotation features to latent regulatory variables in each omic signal, ranked by the relative informativeness in decreasing order. Top 5 most influential genomic annotations are shown in bold for each outlier signal. A detailed explanation of each genomic annotation features included in the model is provided in Supplementary Table 1.

Examining the learned parameters of the multi-omic Watershed model, we observed higher edge weights connecting RNA and methylation, and RNA and splicing signals, compared to those connecting protein and other signals, suggesting varying levels of information sharing between signals in modeling rare variant effects (Figure 4B). Consequently, multi-omic Watershed model outperformed corresponding RIVER models, which were trained on single omic data types at a time, due to information sharing (Figure S12B). The learned weights contributed by each genomic features also reflect known regulatory biology, with distance-based features being highly informative for RNA and methylation outlier signals, splicing annotations most predictive of splicing outlier signal, and missense and loss-of-function annotations most predictive of protein outlier signals (Figure 4C). These results indicate that our multi-omic Watershed model captures biological signals underlying rare variants’ effect on outlier expression across aspects of the regulatory cascade to jointly prioritize functional rare variants.

Given that we observed little difference in the enrichment of rare variant burden when considering all individuals or matching by ancestry within MESA (see Methods), we sought to assess portability of the multi-omic Watershed model across ancestries. We estimated genetic ancestry based on the Human Genome Diversity Panel (HGDP) with seven super populations^13^ and assigned population groups by thresholding the proportion of ancestry estimates (Figure S13). We then trained multi-omic Watershed model using data from N = 426 individuals assigned to European ancestry and evaluated its performance on N2 pair individuals from other populations. We observed comparable predictive performance in terms of area under the precision-recall curve assessment across these populations (Figure S14), suggesting that outlier rare variant effects discovered in one population are likely to exhibit comparable effects across populations, as expected if we are identifying truly causal variants in the absence of significant non-genetic contributions.

### Multi-omics prioritized rare variants are prevalent in each individual

To assess the individual relevance of the rare variants prioritized by the multi-omic Watershed model, we first observed that each individual’s genome had a significant number of rare variants with large posteriors in each omics data type, with 11 RNA variants, 7 splicing variants, 17 methylation variants, and 52 protein variants with posterior >= 0.5 (Figure 5A). Methylation and protein had the highest number of rare variants with predicted large effects at posterior threshold of 0.5 and 0.9. Strikingly, variants prioritized by different outlier signals were largely non-overlapping (Figure S15A), indicating that multi-omic measurements provided complementary information in characterizing effects of rare variants inaccessible to one omics data type alone, as also supported by the increasing enrichment of nearby rare variants when outlier signals are seen at multiple levels.

**Figure 5.**
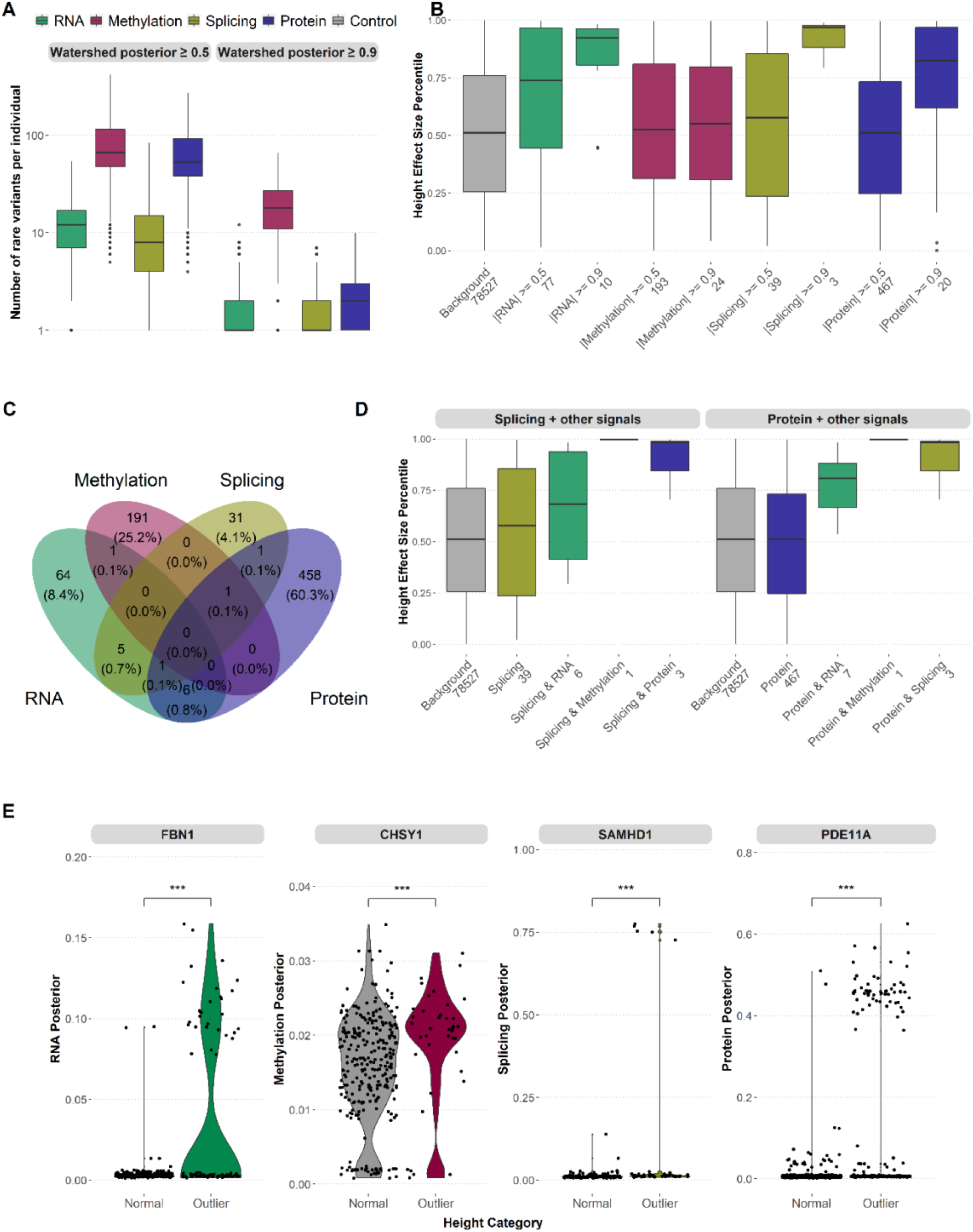
Application of multi-omic Watershed model to inform trait associations. **(A)** Number of rare variants per individual as prioritized by each omic signal (mRNA expression - green; methylation - red; splicing - gold; and protein expression blue) at two levels of Watershed posterior cutoff 0.5 and 0.9. Individuals with significantly large number of outlier expressions (“global outliers”) are removed. The *y*-axis is transformed to log scale. (B) Distribution of percentile normalized effect size for height (median and interquartile range) of all rare variants (background, gray), and those rare variants prioritized by multi-omic Watershed in each signal at two posterior threshold values. Only rare variants mapped to genes with evidence of causing abnormal body height as reported by the Human Phenotype Ontology (HPO) are shown (N = 1,314 genes). Number of rare variants in each category is shown in x-axis labels. Effect size estimate was obtained from UK Biobank GWAS Round 2 using rank-normalized phenotype. (C) Venn diagram of rare variants prioritized by each signal at a posterior threshold of 0.5. (D) Distribution of percentile normalized effect size for height (median and interquartile range) for rare variants prioritized by a single signal at a posterior threshold of 0.5 (left – splicing; right – protein expression) and combined with another signal. (E) Distribution of Watershed posteriors of rare variants identified in individuals with normal height (|Z| < 0.2) and abnormal height (|Z| > 2) in the MESA cohort, collapsed to each gene. Shown are examples of top genes showing differential distribution of posteriors in each omic signal which overlap with HPO annotated genes. *** p < 0.001, one-sided Wilcoxon rank-sum test on absolute value of posteriors between normal and outlier individuals.

To further characterize variants prioritized by each omics data type within an individual, we extracted the probability of loss of function intolerance (pLI) scores^14^ for genes mapped to variants in each group. Notably, pLI scores were not included as an annotation in the Watershed model. We observed that genes with large-effect variants across multiple signals tended to have lower pLI scores and thus were more tolerant of damaging mutations (Figure S15B). Moreover, when we applied MetaDome^15^ to systematically map Watershed-prioritized variants to protein domains, we found that variants falling within homologous meta-domains had higher posteriors in all outlier signals (Figure S15C, p < 2.2e-16). Overall, these data suggest that multi-omic Watershed posteriors captures functional impact of rare variants and provide a strong basis for the application of Watershed posteriors to inform trait associations.

### Multi-omics prioritized rare variants impact multiple complex traits and diseases

We sought to test the hypothesis that rare variants with large multi-omics-based Watershed posteriors are likely to be causal for traits and diseases. We first focused on height, a highly polygenic and heritable trait collected for all individuals in MESA. Based on the summary statistics estimated from a separate cohort, UK Biobank, we identified 78,527 rare variants which overlap with the scored variants in our multi-omic Watershed model which maps to 1,314 genes known to cause abnormal body height as catalogued in the Human Phenotype Ontology (HPO^16^). When restricting to variants prioritized by Watershed with posterior > 0.5 or > 0.9, we observed higher effect sizes on height as compared to background (Figure 5B; Figure S16). The observed higher effect sizes were robust to selecting only the top N variants (N = 10 and 100) from each data type separately (Figure S17A).

Notably, multi-omics outliers could prioritize Mendelian or large-effect genes. We identified a small set of rare variants which were mapped to Mendelian height genes with posterior > 0.5 or were among the top 100 highest posteriors in more than one signal (Figure 5C, S17B). However, when comparing rare variants prioritized by two signals, we found that they had even higher effect sizes compared to those prioritized by single signals, which is especially prominent with splicing and protein when combined with another signal (Figure 5D, Figure S17C-D). Importantly, the shift in effect size by Watershed prioritized variants was also higher than single-omics outliers further stratified by minor allele frequency (MAF, Figure S18), suggesting that the functional correlation between Watershed posterior and effects on trait is not solely driven by MAF. These data support the utility of collecting multi-omic measurements from the same individuals to improve prioritization of functional rare variants with large trait effects that could potentially be missed by traditional approaches such as genome-wide association studies (GWAS).

We next assessed whether Watershed prioritized rare variants could be applied to enhance gene prioritization for complex traits. We obtained body height data on N = 4,559 MESA individuals and after correcting for known covariates, identified those with average body height (“control” individuals, residual |Z| < 0.2) and those with the extremes of body height (“outlier” individuals, residual |Z| > 2). When comparing the distribution of posteriors for all rare variants mapped to each gene between outlier and control individuals, we were able to recapitulate known Mendelian height genes (Figure 5E); importantly, different signals prioritized different genes, which further highlighted the complementary nature of each omic data type. When we combined posteriors across all outlier signals for each variant and compared the resulting gene-level *p*-values with other gene prioritization methods based on common variants (MAGMA^17^), rare coding variants (burden test^18^), or expression quantitative trait loci (PrediXcan^19^), we found that our approach is largely independent (Figure S19), suggesting the unique advantage of incorporating non-coding rare variants in a gene prioritization framework.

To demonstrate the utility of multi-omics prioritized rare variants with Watershed to a diverse range of traits and diseases, we applied similar analyses to immunological and neuropsychiatric traits. Here, we referenced a recent machine learning method that systematically characterized causal genes to primarily focus on rare variants impacting genes with well-predicted trait relevance^20^. For rheumatoid arthritis (RA), we identified 41,339 rare variants with effect size estimates in GWAS and observed that RNA posteriors strongly correlated with effect size (median rank normalized effect size = 0.58 for RNA posterior prioritized RVs compared to 0.50 for background). We further observed that for RA, the protein signal by itself did not prioritize rare variants with large effect size; however, it did when combined with RNA or methylation signals (Figure S20A-B). This observation held true for COVID-19 severity, another immunological trait (Figure S20C-D). For Alzheimer’s disease (AD), methylation outliers were most predictive of effect size, but multimodal outliers have much higher impacts (Figure S21A-C). Interestingly, joint under-expression outliers in RNA and protein signals identified genes with established associations with AD, such as *PDGFRB*^21^ (rs116171826, rs149274963, and rs10071918), *PTN*^22^ (rs61735090), and *MPO*^23^ (rs35897051), supporting the potential role for our prioritized variants in AD pathobiology. For schizophrenia (SCZ), even though we had a smaller set of rare variants with effect size estimates (N=2,851), we observed moderate effect size for variants prioritized by RNA, methylation, and protein signals and a strong shift in splicing prioritized variants (Figure S21D). Notably, in addition to referencing external databases for identifying relevant causal genes, we applied MAGMA to prioritize genes using GWAS summary statistics. We identified 5,378 genes with MAGMA Z > 2 (schizophrenia “positive” genes) and 4,092 genes with MAGMA Z < 0 (schizophrenia “control” genes), and when we compared rare variants with large Watershed posteriors mapped to these two groups of genes, we observed a significant shift in effect size only within positive genes. Overall, these analyses demonstrate that the multi-omic Watershed model represents a flexible framework which can be easily integrated into pipelines for connecting variants to traits.

## Discussion

Rare genetic variants are collectively abundant in the human genome due to recent population expansion^24,25^. They are often population-private, unlike common variants which are shared across populations^26^. Although rare variants have in general larger effects and contribute to risk of complex diseases^27,28^, their abundance may lead to false positive associations and thus require careful methods for analysis and interpretation^29,30^. The present study extends efforts to identify large-effect rare variants through analysis of functional genomics data^1,2,31–35^. By integrating longitudinal multi-omic data collected from a diverse cohort with matched whole genome sequencing, we identified significant enrichment of rare variant burden nearby multi-omic outlier signals across the regulatory cascade.

Our study benefited from both multi-omics data generation and a study design including functional measurements at two time points approximately ten years apart. We observed higher enrichment of rare variant burden in multimodal outliers collected at a single time point compared to joint outliers across two visits based on only a single molecular signal, which indicates that multi-omic datasets can be more beneficial than collecting the same measurement over multiple time points when using those measurements to prioritize functional rare variants.

Importantly, we conducted analyses across an ancestrally-diverse cohort, including individuals with substantial African, East Asian, European, and American (through the inclusion of Hispanic individuals) genetic ancestry, which allows for the evaluation of many additional rare variants than would be included in a cohort containing individuals with all predominantly European ancestry, as is often the case in genomics research due to the over-representation of European populations^36–38^. We found that rare variant enrichments nearby outliers did not change when comparing against all other control individuals as opposed to restricting to controls with similar ancestry, as has been done in previous studies^1,2^. We also found that rare variants associated with multi-tissue expression or splicing changes in GTEx, which consists predominantly of individuals of European ancestry, 15.8% and 15.2% replicated in MESA, which was many more than seen after permuting expression and splicing values. The variants discovered in GTEx that are associated with similar transcriptomic effects in MESA were enriched for rare stop and splice annotations, supporting the use of both genomic annotations and functional signals in variant prioritization.

We extended a Bayesian hierarchical variant effect prediction model, Watershed, to synthesize genomic annotations with observed outlier status in four omics data types. Using multi-omics Watershed, we predicted the functional impact of more than 30 million rare variants and observed that each person in MESA harbors a significant number of rare variants with large posterior probabilities of functional effect. Using this approach, we prioritized multiple novel and known rare variants across common and complex traits and disease including height, Rheumatoid Arthritis, COVID severity, Alzheimer’s Disease and Schizophrenia. Further, we demonstrated how integration of this expanded set of prioritized rare variants aids detection of causal genes.

Combined, we present a comprehensive survey of rare variants underlying multi-omic outlier signals across the regulatory cascade. Using personal multi-omics, our Watershed model prioritized rare variants across a broad range of complex traits. These approaches further demonstrate a general and flexible framework to prioritize impactful rare variants and test for gene associations in diverse population cohorts.

## Online Methods

### The Multi-Ethnic Study of Atherosclerosis (MESA)

MESA is a study of the characteristics of subclinical cardiovascular disease (disease detected non-invasively before it has produced clinical signs and symptoms) and the risk factors that predict progression to clinically overt cardiovascular disease or progression of the subclinical disease^39^. MESA researchers study a diverse, population-based sample of 6,814 asymptomatic men and women aged 45-84. Thirty-eight percent of the recruited participants are white, 28 percent African American, 22 percent Hispanic, and 12 percent Asian, predominantly of Chinese descent. Participants were recruited from six field centers across the United States: Wake Forest University, Columbia University, Johns Hopkins University, University of Minnesota, Northwestern University and University of California - Los Angeles. Participants are being followed for identification and characterization of cardiovascular disease events, including acute myocardial infarction and other forms of coronary heart disease (CHD), stroke, and congestive heart failure; for cardiovascular disease interventions; and for mortality. In addition to the six Field Centers, MESA involves a Coordinating Center, a Central Laboratory, and Central Reading Centers for Computed Tomography (CT), Magnetic Resonance Imaging (MRI), Ultrasound, and Electrocardiography (ECG). The first examination took place over two years, from July 2000 - July 2002. It was followed by five examination periods that were 17-20 months in length. Participants have been contacted every 9 to 12 months throughout the study to assess clinical morbidity and mortality.

Further, the TOPMed MESA Multi-omics Pilot successfully generated transcriptomic data by RNAseq, DNA methylation [850K CpG sites], plasma proteomics by aptamer capture (SomaLogic), and untargeted and targeted metabolomics using liquid chromatography/ mass spectrometry (LC-MS from the Gerszten/Clish laboratory) in ∼1,000 multi-ethnic participants sampled at two time points, Exam 1 and Exam 5, approximately 10 years apart. These data are being used in this study.

### Multi-omic data pre-processing

To identify functional outliers at each time point separately, for three of the four data types considered (total gene expression and splicing measured via RNA-sequencing, DNA methylation, and plasma proteome abundance), we normalized the initial measurements and then corrected for the top 11 genotype principal components, 30 hidden factors determined via PEER^40^, sex, age, and genotype of the strongest QTL per measurement within each exam, before scaling to generate Z-scores on which we threshold to identify outlier events. For assessing DNA methylation outliers, we removed sites if there exists a common SNV that overlap the measurement probe region, and we also removed individual-site pairs if the individual carries a rare variant in the probe’s target region. To estimate methylation at the gene level, we calculate the median methylation Z-score across CpG sites for all sites falling between a gene’s TSS and 1.5kb upstream of TSS^41^. For sOutliers, we define clusters of exon-exon junctions by LeafCutter^42^ from splice junction counts measured by STAR^43^, and identify splicing outlier events using SPOT^2^ which are collapsed to the gene level by taking the minimum *p*-value across clusters per gene.

### Outlier sharing and rare variant enrichment test

We defined outlier (gene, individual) pairs in each omics data to have |Z| > 3 in both time points (joint outliers), where we also assessed outlier sharing across time points and across the regulatory cascade at varying threshold of |Z| (between 2 to 10). For joint outliers in each data type, we calculated mean Z-scores across exams, and assessed outlier sharing by taking a set of (gene, individual) pairs with |Z| > 3 in one signal and calculated proportion of these (gene, individual) pairs in each of the other signals with a relaxed threshold |Z| > 2, for the set of genes and individuals with both data types measured. For expression, methylation, and protein levels, we also assessed outliers with Z < -3 (under outliers) and those with Z > 3 (over outliers) separately. We calculated enrichment as the ratio between outlier and control individuals for the proportion of (gene, individuals) with rare variants within 10kb window of the gene body, restricting to the same set of genes with outlier measurements as before^2^.

### Effects of ancestry on rare variant enrichment

To assess whether the observed enrichments were impacted by differences in genetic ancestry between outlier and non-outlier individuals within each set, even after correcting for the top 11 genotype principal components (PCs) before identifying eOutliers, mOutliers, and pOutliers, we calculated the correlation between individual outlier burden (the number of outliers per data type identified for a single individual) and loadings on genotype PC values for all outlier types, across different thresholds for the definition of outliers. We further replicated rare variant enrichment tests by matching each outlier individual to a non-outlier individual based on closest Euclidean distance defined by the top 11 genotype PC values and compared the resulting relative risk estimates with those from corresponding tests retaining all non-outlier individuals.

### The Multi-omic Watershed model

To integrate genomic annotation with outlier signals to prioritize rare variants with large effects across the regulatory cascade, we extended our Bayesian hierarchical model, Watershed, which consists of a layer of genomic annotation variables (G), a fully connected layer (Z) of latent regulatory variables for each of the four omic signals (mRNA expression, methylation, splicing, and protein expression), and a layer of variables (E) representing the observed outlier status of each omics data type. We used as input p-values for each of the four signals and for all (gene, individual) pairs with at least two signals measured in MESA, along with a set of 77 binary and continuous genomic annotations aggregated across all rare variants nearby each gene. Watershed was then trained to learn edge weights connecting each variable and estimate posterior probability of each rare variant leading to outlier levels of nearby gene for each of the four signals, given genomic annotations and observed expression levels P(E|G, Z). As evaluation, we identified pairs of individuals with the same set of rare variants nearby the same gene (N2 pair), and asked the Watershed model to predict the regulatory status of one individual in the pair based on genomic annotations of rare variants and observed outlier status of each omic signal in the other individual. We benchmarked the performance of multi-omic Watershed model against logistic regression models trained using genomic annotations alone (GAM) and Bayesian models based on single outlier signals at a time (RIVER), at the same p-value threshold for defining outliers (p < 0.05). We repeated the same analysis when training the model on individuals of European ancestry and evaluating its performance on individuals from other ancestries to assess its cross-population portability.

### Application of multi-omic Watershed posteriors to trait analysis

To test the hypothesis that rare variants with large Watershed posteriors are likely to be causal for traits through altered expression of nearby genes, we compared Watershed posterior probabilities against variant effect sizes estimated from genome-wide association studies (GWAS) obtained from independent datasets. For each trait, we considered the set of rare variants reported in GWAS summary statistics which were also present in MESA and applied rank normalization on effect size for all such rare variants (background). The distribution of normalized effect size was then compared across rare variants prioritized by Watershed at varying thresholds (0.5 or 0.9), where we focused on variants nearby genes with known evidence of being causal for each trait through the Human Phenotype Ontology (HPO^16^) or Open Targets^20^. Given the differences in posterior distribution in each omics data type, we also considered same number of top N variants as prioritized by Watershed for each data type in isolation (N = 10 and 100) and compared their effect size distribution against the background. Additionally, because Watershed can leverage data from all outlier signals and make posterior predictions even for unobserved omic measurements, we repeated this analysis after subsetting to variants mapped to directly measured genes in each signal.

To assess whether Watershed posteriors can be applied to prioritize genes, we adopted a similar approach as rare variant collapsing analysis by aggregating all rare variants within 10kb window of a gene and compared the distribution of rare variant posteriors for individuals at the extremes of trait. Specifically for height, we first calculated residual height after regressing out age, sex, self-reported race, clinical center, and top 10 genotype PCs, for N = 4,559 individuals in MESA for whom we collected height and demographic data. We then defined individuals with residual |Z| > 2 as having outlier height and those with residual |Z| < 0.2 as controls. For each gene, we compared Watershed posterior distribution for all rare variants in outlier individuals versus controls using a Wilcoxon rank-sum test and controlled for multiple testing by the Benjamini-Hochberg procedure.

## Acknowledgments

The authors would like to thank V. Wang, J. Weinstock, and T. Nguyen for helping with data management, and members of the Montgomery and Battle labs for input on interpretation of results. AB, NS, JB, and DA were supported by R01HL141989. NMF was supported by the National Science Foundation Graduate Research Fellowship, grant no. DGE – 1656518 and a graduate fellowship from the Stanford Center for Computational, Evolutionary and Human Genomics. TL was supported by T32 GM136577. SBM was supported by R01AG066490, R01MH125244, U01HG009431 (ENCODE) and R01HL142015 (TOPMed).

Whole genome sequencing (WGS) for the Trans-Omics in Precision Medicine (TOPMed) program was supported by the National Heart, Lung and Blood Institute (NHLBI). WGS for “NHLBI TOPMed: Multi-Ethnic Study of Atherosclerosis (MESA)” (phs001416.v1.p1) was performed at the Broad Institute of MIT and Harvard (3U54HG003067-13S1). Centralized read mapping and genotype calling, along with variant quality metrics and filtering were provided by the TOPMed Informatics Research Center (3R01HL-117626-02S1). Phenotype harmonization, data management, sample-identity QC, and general study coordination, were provided by the TOPMed Data Coordinating Center (3R01HL-120393-02S1), and TOPMed MESA Multi-Omics (HHSN2682015000031/HSN26800004). The MESA projects are conducted and supported by the National Heart, Lung, and Blood Institute (NHLBI) in collaboration with MESA investigators. Support for the Multi-Ethnic Study of Atherosclerosis (MESA) projects are conducted and supported by the National Heart, Lung, and Blood Institute (NHLBI) in collaboration with MESA investigators. Support for MESA is provided by contracts 75N92020D00001, HHSN268201500003I, N01-HC-95159, 75N92020D00005, N01-HC-95160, 75N92020D00002, N01-HC-95161, 75N92020D00003, N01-HC-95162, 75N92020D00006, N01-HC-95163, 75N92020D00004, N01-HC-95164, 75N92020D00007, N01-HC-95165, N01-HC-95166, N01-HC-95167, N01-HC-95168, N01-HC-95169, UL1-TR-000040, UL1-TR-001079, UL1-TR-001420, UL1TR001881, and DK063491. The authors thank the other investigators, the staff, and the participants of the MESA study for their valuable contributions. A fill list of participating MESA investigators and institutes can be found at http://www.mesa-nhlbi.org.

