## Supplementary Information for "The functional impact of rare variation across the regulatory cascade"

### Methods

*MESA study*

The Multi-Ethnic Study of Atherosclerosis (MESA)[1] was designed as a study of the characteristics of subclinical cardiovascular disease and the risk factors that predict disease progression. MESA recruited a sample of 6,814 asymptomatic men and women aged 45-84, and specifically sought to obtain a diverse population sample, resulting in 38% white participants, 28% African-American, 22% Hispanic, and 12% Asian, predominantly of Chinese descent. Participants were recruited from six locations, including Wake Forest University, Columbia University, Johns Hopkins University, University of Minnesota, Northwestern University and University of California at Los Angeles. We retain only unrelated samples across all analyses.

*RNA-sequencing data processing*

The scripts and reference annotations used to quantify transcripts mapping to each gene are available and described here: https://github.com/broadinstitute/gtex-pipeline/blob/master/TOPMed_RNAseq_pipeline.md. Briefly, RNA-seq reads were aligned using STAR[2] to the GRCh38 reference genome, and gene quantification and quality control was done using RNA-SeQC[3], resulting in read counts and number of transcripts per million mapped reads (TPM). As described in chapter 3, we log2-transformed the expression values ($\log_2$(TPM + 2)), using the GENCODE v30 gene annotation, available at the above URL. We subsetted to autosomal lincRNA and protein-coding genes and restricted to genes with at least 6 reads and TPM > 0.1 in at least 20% of individuals.

*DNA methylation data processing*

DNA methylation measurements were obtained from whole blood using the Illumina EPIC chip. Initial quality control and normalization was performed using the meffil R package[4] for functional normalization. Briefly, quality control included assessing sample swaps, of which two were identified and resolved, sample call rate, sex detection mismatches, and genotype concordance. We exclude samples where >5% of CpG sites had a detection p-value > 0.01 and those that were visible outliers based on their ratio of methylated to unmethylated signal. We also remove three samples whose genotypes did not match the SNP probes included on the microarray (concordance threshold = 0.8). After removing samples that failed QC, we apply functional normalization[5], which extends the idea of quantile normalization to adjust for unwanted technical variation via control probe PCs. We used 10-fold cross validation to determine the number of control PCs to include based on the residual variance after fitting 20 PCs, and decided to include 11, based on decreases in the residuals. We apply some additional filtering: (1) Remove individual-site instance if detection p-value > 0.01, (2) Exclude probes if > 10% of values are missing, (3) Several filters based on mapping issues, non-CpG targeting, or polymorphisms as described in[6], with suggested masking and variant annotations for this array available here: http://zwdzwd.github.io/InfiniumAnnotation.

*Plasma proteome data processing*

Proteome measurements were obtained via the SOMAscan HTS Assay 1.3K - Plasma, which is a highly multiplexed, aptamer-based assay[7], with assessment of this array in particular described in[7,8]. SomaLogic suggests several normalization steps, starting with the observed Relative Fluorescence Intensities (RFUs) which are compared against reference values and scaled accordingly (denoted by SomaLogic as Hybridization Control Normalization or Hyb). They also apply Median Signal Normalization (Hyb.MedNorm), which is an intraplate normalization procedure to remove sample-to-sample differences that may be due to overall protein concentration or experimental variation. Then there is a calibration step (Hyb.MedNorm.Cal), based on the levels of each analyte within calibrator replicates. These steps are described in further detail in[8].

*Variant calling and annotations*

Whole genome sequencing was generated as described in[9]. We restrict our analysis to single nucleotide variants and small insertions and deletions that appear at a less than 1% frequency across MESA as well as < 1% across the entire gnomAD dataset[10] and < 1% in all relevant gnomAD sub-populations, including non-Finnish European, African, East Asian, and American. We annotated the VCF using Ensembl VEP (version 103[11]). CADD[12] scores were extracted from a pre-compiled annotation file (https://cadd.gs.washington.edu/download) using variant scores from the hg38 genome build.

*eOutlier calling*

To identify expression outliers, we take the log-transformed TPM values within each exam for autosomal lincRNA and protein-coding genes and identify 30 hidden factors associated with technical variation via PEER[13]. We then run a linear regression model with the log-transformed TPM values as the outcome variable and the 30 hidden factors, top 11 genotype PCs, genotype of the strongest cis-eQTL per gene, age, and sex as predictors.

We compute the model residuals using the lm() function in R. We scale the residuals from that regression to generate Z-scores within each exam. We then require either the outlier or non-outlier signal at a given threshold to be seen in both exams for all downstream analyses, except for assessing replication across exams, where we apply the thresholds in each exam separately. We remove individuals that have a number of outlier genes ($|Z| > 3$ in both exams) greater than 1.5*IQR based on the distribution of eOutlier burden across all individuals.

*mOutlier calling*

To identify methylation outliers, we first transform the normalized beta values to m-values[14]:

$$m = \log_2(beta_i / (1 - beta_i))$$

From there, we apply the same correction approach as described above, though hidden factors via PEER[13] were learned on inverse normalized beta values from a random subset of 100,000 CpG sites. We correct for 30 PEER factors, the top 11 genotype PCs, genotype of the strongest cis-mQTL per gene (with FDR < 0.25), age, and sex, and again scale the residuals to generate Z-scores per CpG site. We then filter out sites with any common SNVs overlapping the measurement probe, including the CpG site, and individual-site instances if the individual carries a rare variant within the probe region, and retain only autosomal sites. We remove individuals that have a number of outlier CpG sites (|Z| > 3 in both exams) greater than 1.5*IQR based on the distribution of mOutlier burden across all individuals. To calculate gene-level methylation Z-scores, we take the median Z-score across all measured CpG sites within 1.5kb upstream of a gene's TSS, restricting to protein-coding and lincRNA genes, as in the expression analyses.

*sOutlier calling*

We applied the SPOT (SPlicing Outlier deTection) framework to detect splicing outliers similar to previous work[15]. Briefly, we performed intron clustering by adapting a LeafCutter pipeline[16] from STAR-aligned junction reads. We applied custom filtering to remove exon-exon junctions with low expression while retaining rare junctions by excluding junctions where no sample has >= 15 reads, and further excluded exon-exon junctions with less than 40% of samples with more than three reads. We then applied the SPOT pipeline to first fit a Dirichlet-Multinomial distribution to counts spanning alternatively spliced exon-exon junctions for each gene, based on which we then identified individuals with significant deviation away from the population mean based on Mahalanobis distance (MD) metric. To avoid biases caused by dimensionality where clusters with larger number of exon-exon junctions show different MD distribution compared to smaller clusters, we computed the empirical p-values for each individual in each cluster by comparing against 1 million random samples from the fitted Dirichlet-Multinomial distribution. To map empirical p-values from intron clusters to genes, we took the minimum p-value ($p_m$) across all c clusters within the span of each gene and computed the conservative estimate of the probability of observing pm across c independently drawn uniform distributions as

$$P_{gene} = 1 - (1 - p_m)^c$$

Finally, we converted p-values to z-scores assuming a normal distribution.

*pOutlier calling*

For protein outliers, we natural log transform the normalized fluorescence values, before proceeding with the same correction procedure, where we identify 30 hidden factors via PEER[13] and correct for those in addition to the top 11 genotype PCs, genotype of the strongest cis-pQTL per gene (with FDR < 0.25), age, and sex, before scaling the residuals within each exam to generate Z-scores. We remove individuals that have a number of outlier proteins (|Z| > 3 in both exams) greater than 1.5*IQR based on the distribution of pOutlier burden across all individuals.

*Enrichments*

Enrichments were calculated by restricting SNVs and indels to those that occur at less than 1% frequency across MESA and for those found in gnomAD[10], also at less than 1% frequency across all of gnomAD, as well as relevant sub-populations (see above). For eOutliers and pOutliers, we intersect variants with the gene body +\- 10kb on either end of the gene, based on gencode v26 annotations: https://www.gencodegenes.org/human/release_26.html. For mOutliers, we intersect variants with varying window sizes around the CpG site, using bedtools[17]. After intersecting variants, we convert to a binary signal, with 1 indicating at least one rare variant was found in the region for that individual and 0 indicating no rare variants. For additional annotations, we subset the full set of rare variants to those with the given annotation, as determined via VEP[11], release 103, and CADD[12], version 1.6. Additionally, we assessed whether each variant lies within protein meta-domains as identified by MetaDome[18] after lifting over all variants to hg19.

We calculate relative risk as the proportion of outliers with a nearby rare variant divided by the proportion of non-outliers with a nearby rare variant, restricting the set of non-outliers to the same set of genes for which outlier individuals were discovered. We use the epitools R package to estimate the relative risk and confidence intervals. We define non-outliers as those with $|Z| < 1$ in both exams.

*Matching controls by genotype PC*

To assess whether observed enrichments are due to differences in genetic ancestry between outlier and control individuals, we also calculate enrichments where we select a control individual for each outlier by taking the individual from the full set of controls (|exam 1 Z-score| < 1 and |exam 5 Z-score| < 1) with the lowest Euclidean distance to the outlier individual based on the top 11 genotype PC values. Distances were calculated using the philentropy R package.

*GTEx replication*

We subset rare variants seen in GTEx to those associated to multi-tissue expression outlier effects. We assess the subset of the rare SNVs and indels observed in GTEx that were also seen in any individuals in MESA and calculate the proportion that also lead to observable outlier effects at the methylation, expression, and protein level. We assess significance of this overlap based on permutations, where within MESA, we permute Z-scores, keeping measurements together across time points, within each gene across individuals and assess the number of times a GTEx eOutlier variant is associated with an expression change in MESA.

To assess enrichment of different types of variants in the set of replicating variants as compared to the remaining variants, for each annotation, we create a contingency table where the rows indicate whether or not the variant effect from GTEx was observed in MESA and the columns contain the number of variants with and without the given annotation. We then calculate a relative risk of a variant associated with a replicating effect having a given predicted effect, i.e. annotation again using the epitools R package to estimate the relative risk and confidence intervals.

*The multi-omic Watershed model*

Watershed was designed to model instances of (gene, individual) pairs given the functional annotations of nearby rare variants and observed outlier status of gene measurements. The multi-omic Watershed model requires two kinds of input variables:

1. A set of genomic annotation variables for each rare variant: We curated a list of N = 73 annotations for each variant, consisting of variant effect predictor (VEP) consequences, regulatory element annotations, conservation scores, and other genomic and epigenomic features from other models such as CADD and ENCODE, detailed in Supplementary Table 1. For each (gene, individual) pair, we aggregated each annotation across all rare variants within the 10kb window of the gene. Finally, we performed mean centering and scaling for each genomic annotation variable as input to the Watershed and GAM models.
2. A set of categorical variables representing the observed outlier status of the gene: Based on the estimated p-values in each signal (mRNA expression, methylation, splicing, and protein expression), we binarized our (gene, individual) pairs as outliers in each dimension at a *p*-value threshold of 0.05 (--pvalue_threshold=0.05 as input to the model). In practice, this translates roughly to the top 2% of Z-scores in each signal (at a Z-score cutoff of around 2), which we also used for evaluation of the model (--pvalue_fraction=0.02). We explicitly modeled over- and under-outliers for RNA, methylation, and protein levels.

To enrich for outlier signals with genetic effects, we only included genes with consistent Z-scores across the two visits by removing (gene, individual) pairs where one visit has |Z| >=3 and the other has |Z| <= 1. We used median Z-scores across visits as input to Watershed. We also removed individuals who have significantly higher number of outlier measurements (“global outliers”), defined as those having more than Q3 + 1.5*IQR outliers in each signal (Q3 represents 75th percentile rank value and IQR represents interquartile range). Further, to model relationships between omic signals, we only kept (gene, individual) pairs with at least two types of omic measurements (out of four), and filtered out genes which do not have any outlier individual in at least two measurements.

We applied Watershed exact inference optimization routine which is tractable for K = 4 outlier signals. To evaluate the performance of the multi-omic Watershed model, we took pairs of individuals with the same set of rare variants nearby the same gene (“N2 pairs”), who were not included in the training of the model, to assess the ability of the model to predict regulatory status of the second individual based on genomic annotations and observed outlier status of the first individual in each dimension, using area under the precision-recall curve as a metric. In total, we had 596,288 instances of (gene, individual) pairs, of which we had 60,724 N2 pair individuals for evaluation of the model.

To estimate Watershed posterior probabilities, we applied the trained multi-omic Watershed model and calculated P(Z | G, E) for all rare variants from all (gene, individual) pairs in MESA (description of variables in Figure S12A). We scored posterior probability for a total of 30 million (gene, individual, rare variant) triplets. Importantly, Watershed can model missing outlier measurements such that each variant has posterior calculated in all four signals. We assigned a final posterior estimate for each rare variant by taking the maximum across all individuals with the variant.

*GAM and RIVER*

We followed previous procedures in training genomic annotation models (GAM) and RIVER models[19]. Briefly, for GAM, we applied a logistic regression model using all genomic annotations as input features and l2 regularization to promote sparsity. We used as the response variable the binarized outlier status (a *p*-value threshold of 0.05) for each signal. For RIVER, we ran separate Watershed models with one outlier signal in each because it is a special case of Watershed.

*Cross-population comparison of Watershed performance*

Based on the seven super populations present in the Human Genome Diversity reference panel[20], we applied RFmix[21] to estimate genetic ancestry for each individual in MESA. We assigned individuals to ancestries if they have a probability > 0.75 of belonging to the group, with the exception of Hispanic population (Native Americans) where we used a threshold of 0.5 to include more people. As a result, we identified 426 Europeans, 270 Africans, 107 East Asians, and 54 Hispanic individuals.

To assess cross-population portability of multi-omic Watershed, we constructed training data from 147,569 (gene, individual) pairs, following the same inclusion and exclusion criteria as the main model, where all individuals are of European ancestry. We then constructed evaluation data using N2 pairs where both individuals come from the same population. We applied the Watershed model trained from European individuals and tested its performance separately on N2 pairs from other populations in each omic signal, where we excluded populations with less than ten N2 pairs in each signal for evaluation. To compare statistical difference, we designed a bootstrapping procedure to subsample half of N2 pairs in each test set 100 times and computed the distribution of area under the precision-recall curve across bootstrapped samples.

*Correlation of Watershed posteriors with GWAS effect size*

Because Watershed predicts functional impact or rare variants across the regulatory cascade, we reasoned that Watershed prioritized variants affecting essential genes related to a trait should have a high functional impact on the trait. To systematically test this hypothesis, we identified several polygenic traits with summary statistics and compared the distribution of estimated effect size with variants prioritized by Watershed at different posterior thresholds.

Specifically, we considered the following traits：

- Height: We obtained summary statistics on rank-normalized standing height from UK Biobank release 2 with 361,194 individuals of both sexes. This data includes imputed genotypes from HRC plus UK10K & 1000 Genomes reference panels as released by UK Biobank in March 2018.

- Rheumatoid arthritis: We obtained summary statistics from a trans-ethnic meta-analysis with 19,234 cases and 61,565 controls[22].
- COVID-19 severity: We obtained summary statistics comparing severe positive cases with non-severe positive cases based on 1,244 cases and 16,413 controls from UK Biobank released in May 2021.
- Alzheimer's Disease (AD): We obtained summary statistics from a recent meta-analysis which includes 90,338 cases and 1,036,225 controls[23].
- Schizophrenia: We obtained summary statistics from the most recent public release from the Psychiatric Genomics Consortium (wave 3) consisting of 67,390 cases and 94,015 controls[24].

For each trait, we first lifted the summary statistics to GRCh38 genome assembly and intersected with all rare variants present in MESA. This step resulted in varying amount of variants left because of differences in sequencing platform and imputation strategy from different studies. In total, we had a range of thousands (schizophrenia) to hundreds of thousands (COVID-19 severity) rare variants with both effect size estimation and posterior probabilities leading to outlier levels in four omic signals from multi-omic Watershed. We then applied percentile normalization on variant effect sizes and compared distribution of normalized effect size of Watershed prioritized rare variants to those of the background (all rare variants regardless of Watershed posterior). We set threshold of posterior probabilities in each signal to prioritize variants; however, due to the difference in the distribution of posterior probabilities across outlier signals (e.g. methylation and protein expression have larger number of high-posterior variants than RNA expression and splicing), we also chose top N variants ranked by posteriors in each comparison. Importantly, to enrich for functional variants most likely to affect traits, we focused on genes with evidence of association with each traits using either known associations from the Human Phenotype Ontology (HPO[25]) and OpenTargets[26], or applied MAGMA gene-level tests[27] to define genes most relevant for the trait.

*Gene-based test for association with height using Watershed posteriors*

Among the 1,319 individuals with multi-omic measurements in MESA, we computed posterior probabilities for each rare variant leading to outlier RNA expression, methylation, splicing, or protein expression in nearby genes. We reasoned that these posteriors can directly reflect the functional impact of these variants on traits, particularly if they map to genes with strong evidence of association with the trait. We also noted that once trained, these posteriors can be applied to a much larger set of individuals with whole genome sequencing data for association testing as long as they share those rare variants identified in the original training cohort; therefore, Watershed posteriors can be used in a general framework as weights for gene association testing, most appealingly in biobank scale data which do not need to have multi-omic measurements.

As a proof of concept for this workflow, we collected standing height from 4,559 individuals with whole genome sequencing and computed residual height after regressing out age, sex, self-reported race, clinical center, and top 10 genotype PCs. We defined individuals with residual

|Z| > 2 as outlier individuals, and those with residual |Z| < 0.2 as controls, resulting in 182 outlier individuals and 680 control individuals. We next collapsed all rare variants located within 10kb of the gene body for each gene, and applied Wilcoxon rank-sum tests to assess the difference in distribution of posteriors in outlier and control individuals. This gene-level test is similar to the burden test framework but uses Watershed posteriors as weights and incorporates many more non-coding variants. We applied this analysis using posteriors from each signal and a combined posterior summarizing the largest effects across all four signals.

We compared our test with three well-established gene prioritization methods：

1. MAGMA: we applied MAGMA gene analysis[27] using the same GWAS summary statistics on height from UK Biobank and the top SNP model to derive gene-level p-values.
2. PrediXcan: we applied PrediXcan using the same GWAS summary statistics on height from UK Biobank (S-PrediXcan) and MASHR-based eQTL models trained across 49 tissues in GTEx[28–30]. We chose the minimum p-value across tissues to obtain gene-level *p*-values.
3. Burden test: we obtained precalculated burden test results from Helix[31] which is also based on UK Biobank data. We used BOLT-LMM p-values in our comparison.

*Data Availability*

CADD variant annotations: https://cadd.gs.washington.edu/download
VEP variant annotations: https://useast.ensembl.org/info/docs/tools/vep/index.html
Human phenotype ontology (HPO): https://hpo.jax.org/app/
Open Targets: https://www.opentargets.org/
UK Biobank GWAS summary statistics: http://www.nealelab.is/uk-biobank
Psychiatric Genomics Consortium (PGC) GWAS summary statistics: https://www.med.unc.edu/pgc/download-results/
Rheumatoid arthritis GWAS summary statistics: http://plaza.umin.ac.jp/~yokada/datasource/software.htm
COVID-19 severity summary statistics: https://grasp.nhlbi.nih.gov/Covid19GWASResults.aspx
Burden test data from Helix https://s3.amazonaws.com/helix-research-public/ukbb_exome_analysis_results/README.txt

*Code Availability*

Watershed model: https://github.com/BennyStrobes/Watershed
MAGMA software https://ctg.cncr.nl/software/magma
PrediXcan software https://github.com/hakyimlab/MetaXcan

### Supplemental Figures

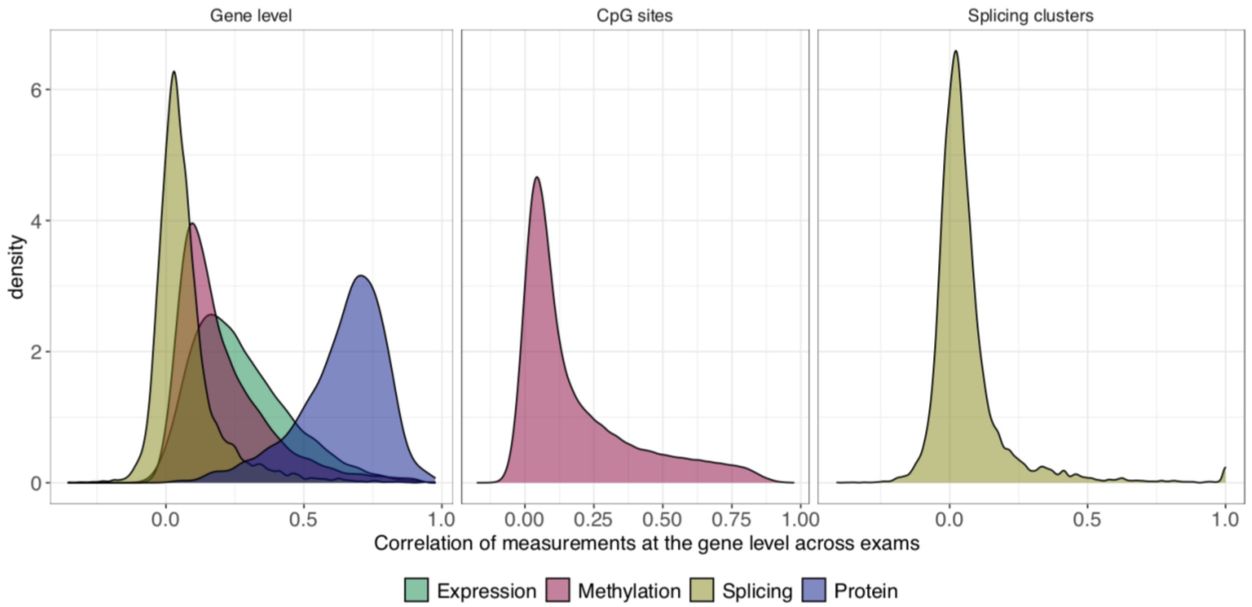


**Figure S1. Correlation of corrected measurements over time.** The distribution of Pearson correlation coefficients across expression Z-scores (green), gene-level methylation Z-scores (red), gene-level splicing Z-scores (gold), and protein Z-scores (blue) on the left, with the correlation across Z-scores per CpG sites in the center and across splicing clusters on the right.

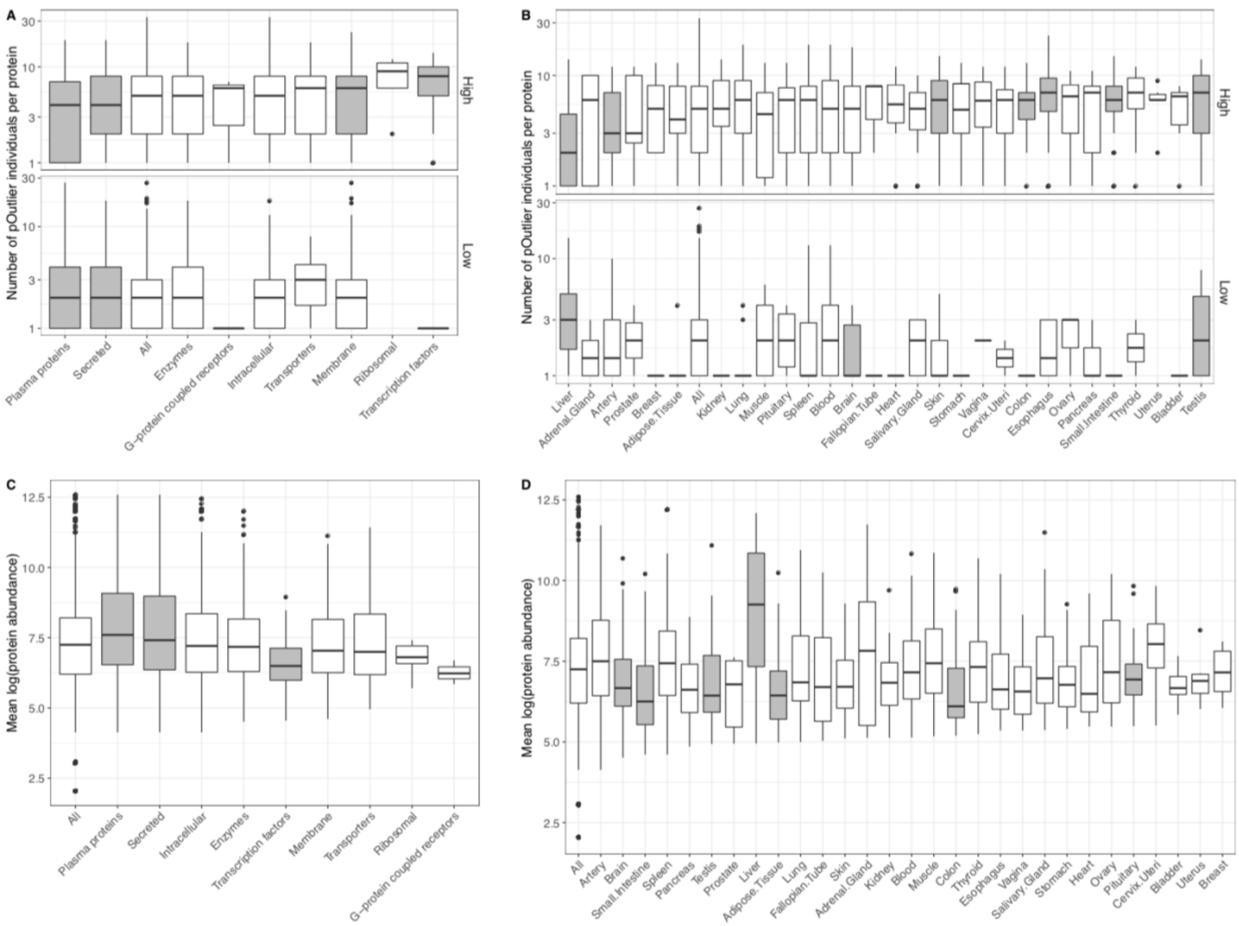


**Figure S2. Distribution of pOutlier burden across proteins by class and tissue. (A)** The distribution of number of pOutlier individuals across proteins within different classes, as annotated by the Human Protein Atlas (*16*), split by the direction of effect with high abundance pOutlier burden on the top and low abundance pOutlier burden on the bottom. **(B)** The mean log(protein abundance) across all individuals for proteins annotated to different classes. **(C)** The distribution of number of pOutlier individuals across proteins annotated as either enhanced or enriched for given GTEx tissues, split by the direction of effect with high abundance pOutlier burden on the top and low abundance pOutlier burden on the bottom. **(D)** The mean log(protein abundance) across all individuals for proteins annotated to different tissues. For all panels, the shaded grey boxes indicate a significant difference between the values for that class or tissue and the set of all proteins, as determined by a two-sided Wilcoxon rank sum test with $p < 0.05$.

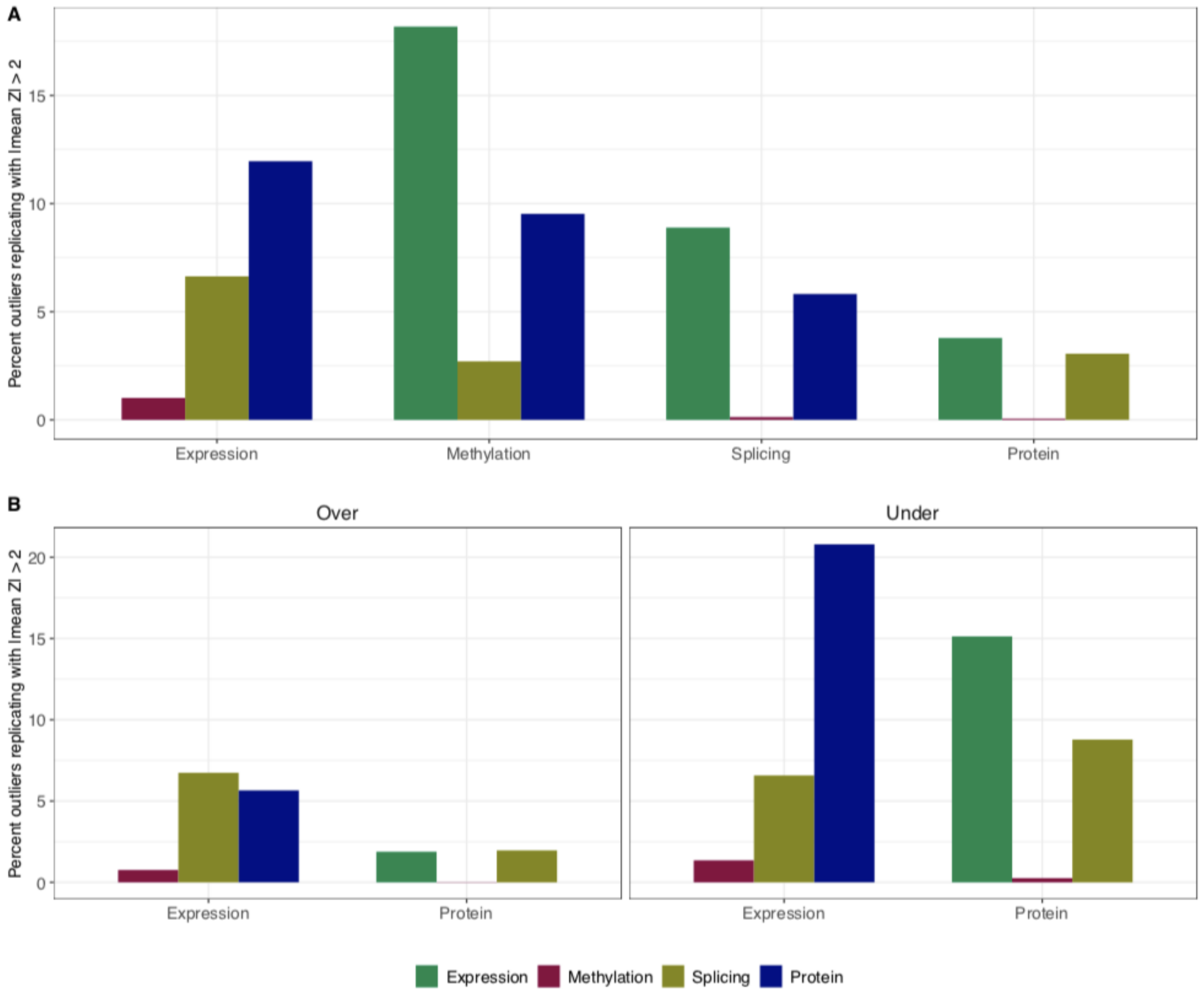


**Figure S3. Replication of outliers across data types. (A)** The percent of outliers (y-axis) identified within each data type (x-axis) that are also seen across each other data type at a threshold of |mean Z| > 2 across exams, considering the set of genes and individuals measured in both. **(B)** The percent of eOutliers and pOutliers (y-axis) identified within each data type (x-axis) that are also seen across each other data type at a threshold of |mean Z| > 2 across exams, considering the set of genes and individuals measured in both, split by the direction of the outlier effect in the discovery data type.

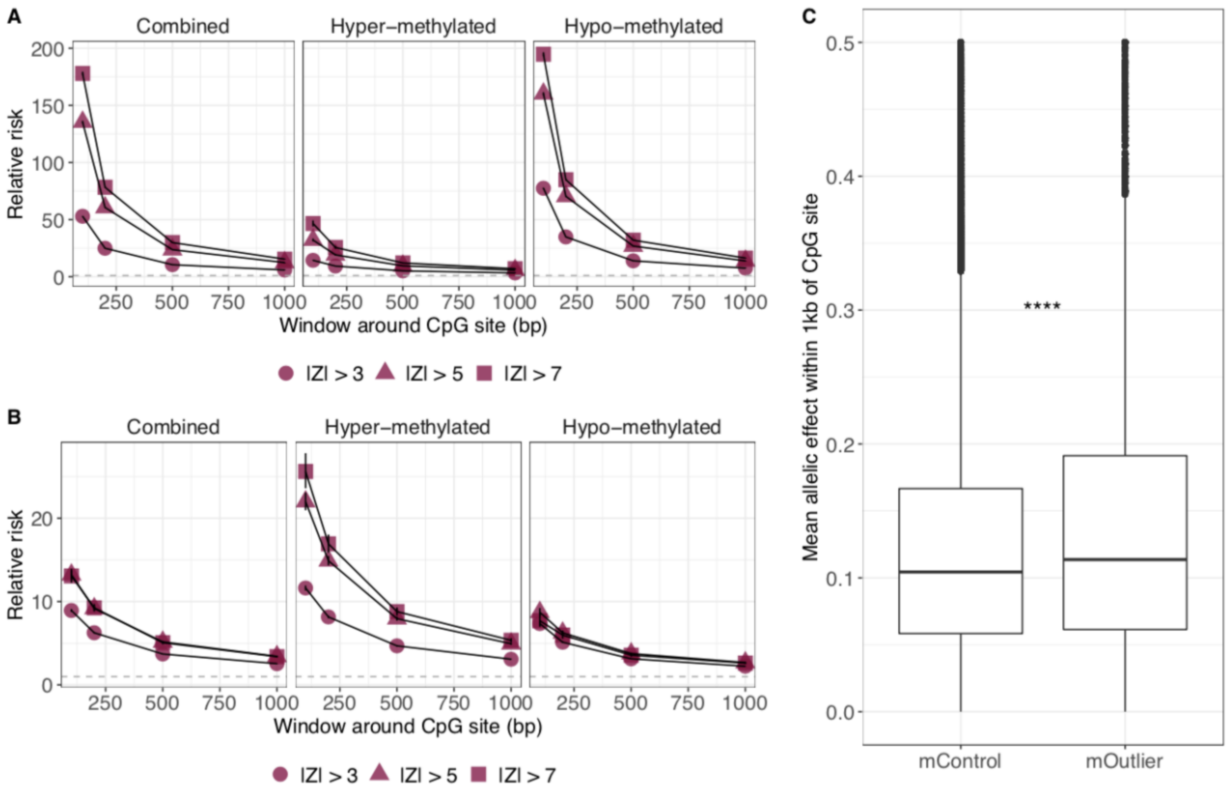


**Figure S4. Enrichment of rare variants nearby CpG-level mOutliers. (A)** The relative risk of carrying a nearby rare variant within varying distances from the outlier CpG site (x-axis) across thresholds (shapes) for CpG-level mOutliers identified across both exams (left), as well as the subset that show hyper-methylation (center) and hypo-methylation (right). Any probes that overlap common SNVs have been filtered out, as well as individual-probe instances if a rare variant overlaps the measurement probe as well. **(B)** The relative risk of carrying a nearby rare variant within varying distances from the outlier CpG site (x-axis) across thresholds (shapes) for CpG-level mOutliers identified across both exams (left), as well as the subset that show hyper-methylation (center) and hypo-methylation (right). Any probes that overlap common SNVs have been filtered out, as well as individual-probe instances if a rare variant overlaps the measurement probe and/or the CpG site as well. **(C)** For the set of CpG-level mOutliers assessed in (A), the mean allelic effect measured for any SNVs within a 1kb window around mOutliers as well as control individuals for the same set of sites. $p$ < 2.2e-16 based on a one-sided Wilcoxon rank sum test.

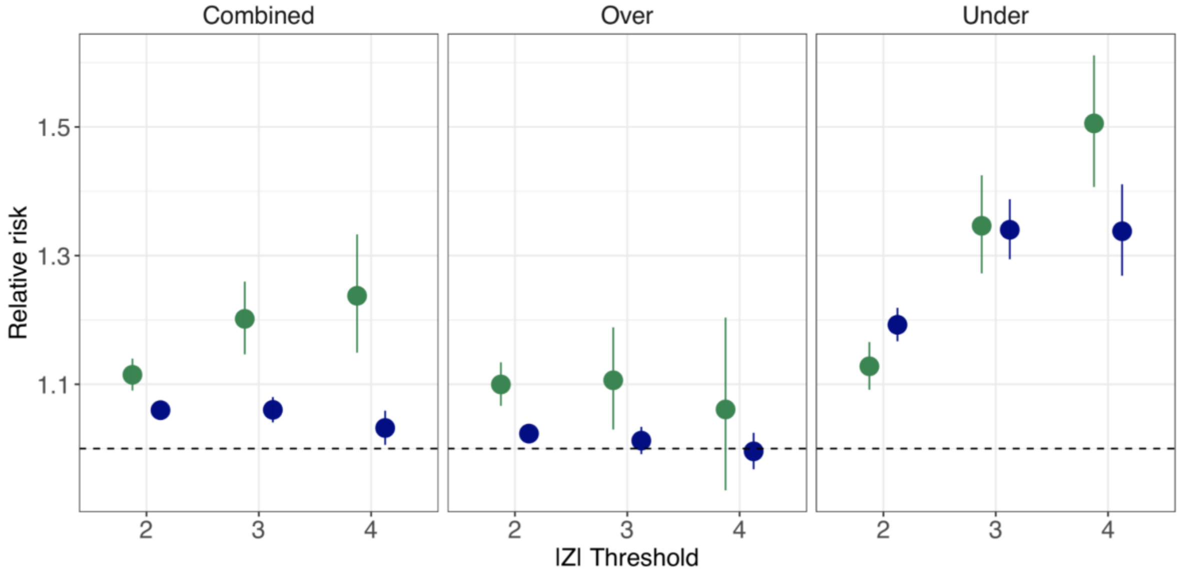


**Figure S5. Enrichment of eOutliers restricted to the set of assayed protein genes.** Estimates of relative risk of carrying a nearby rare variant (gene body +/- 10kb) across thresholds (x-axis) for eOutliers (green) identified across both exams for the subset of genes with protein measurements, and all pOutliers plotted for comparison (blue).

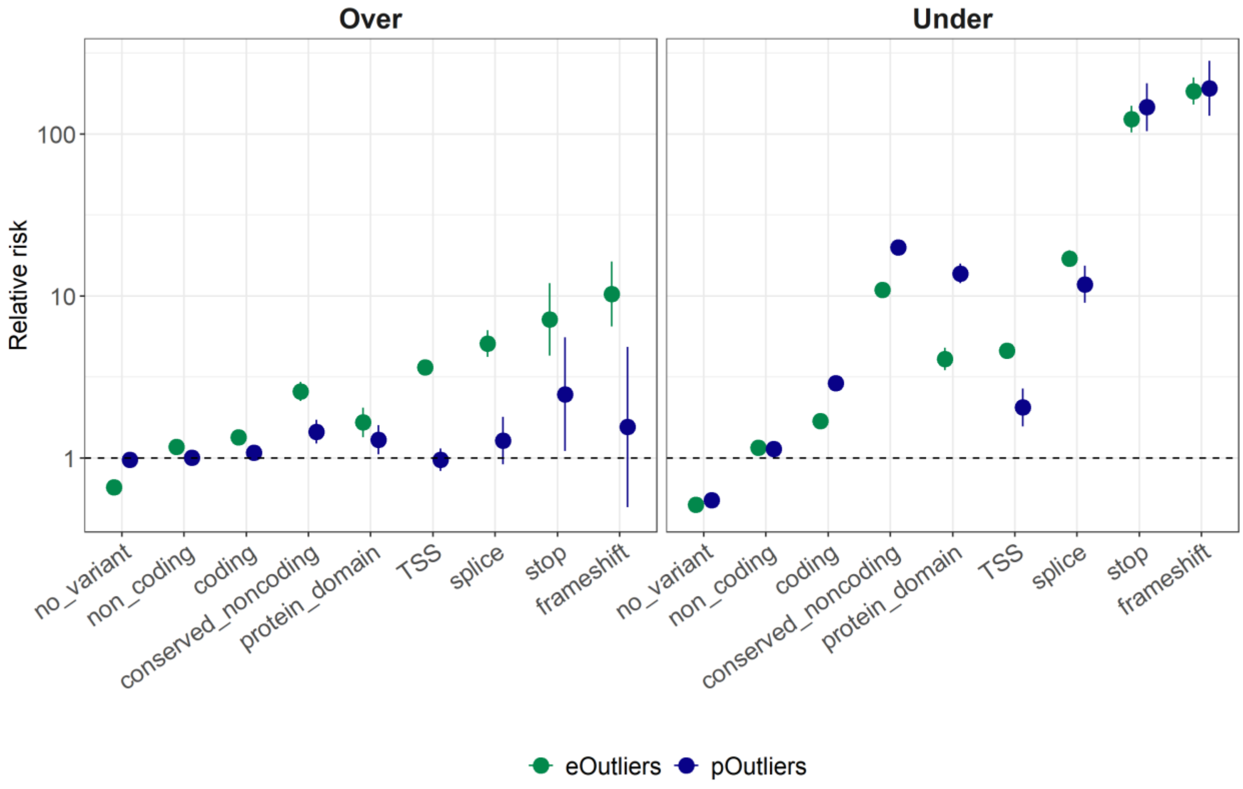

**Figure S6. Enrichment of annotated rare variants nearby eOutliers and pOutliers.** Estimates of relative risk of carrying a nearby rare variant (gene body +/- 10kb) for different types of variants (x-axis) for eOutliers (green) and pOutliers (blue) split by the direction of the effect.

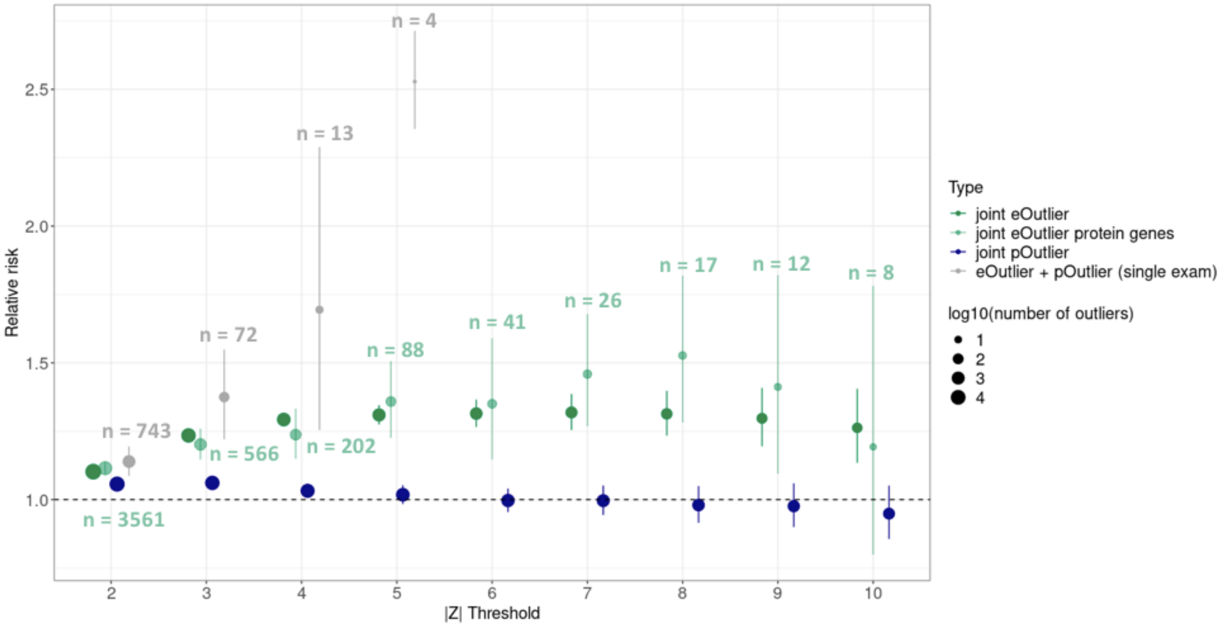


**Figure S7. Enrichment of single time point overlapping outliers vs joint outliers in a single data type. (A)** The relative risk of carrying a nearby rare variant (gene body +/- 10kb) across thresholds (x-axis) for eOutliers identified across both exams (green), the subset of joint eOutliers for genes also measured in protein (light green), pOutliers identified across both exams (blue), and overlapping eOutliers and pOutliers identified in a single exam (grey). The number of identified outliers are listed at each threshold for the filtered joint eOutlier set (light green) and the overlapping outlier set (grey), with the size of each point also varying by the log10(number of outliers) seen for that particular set and threshold combination.

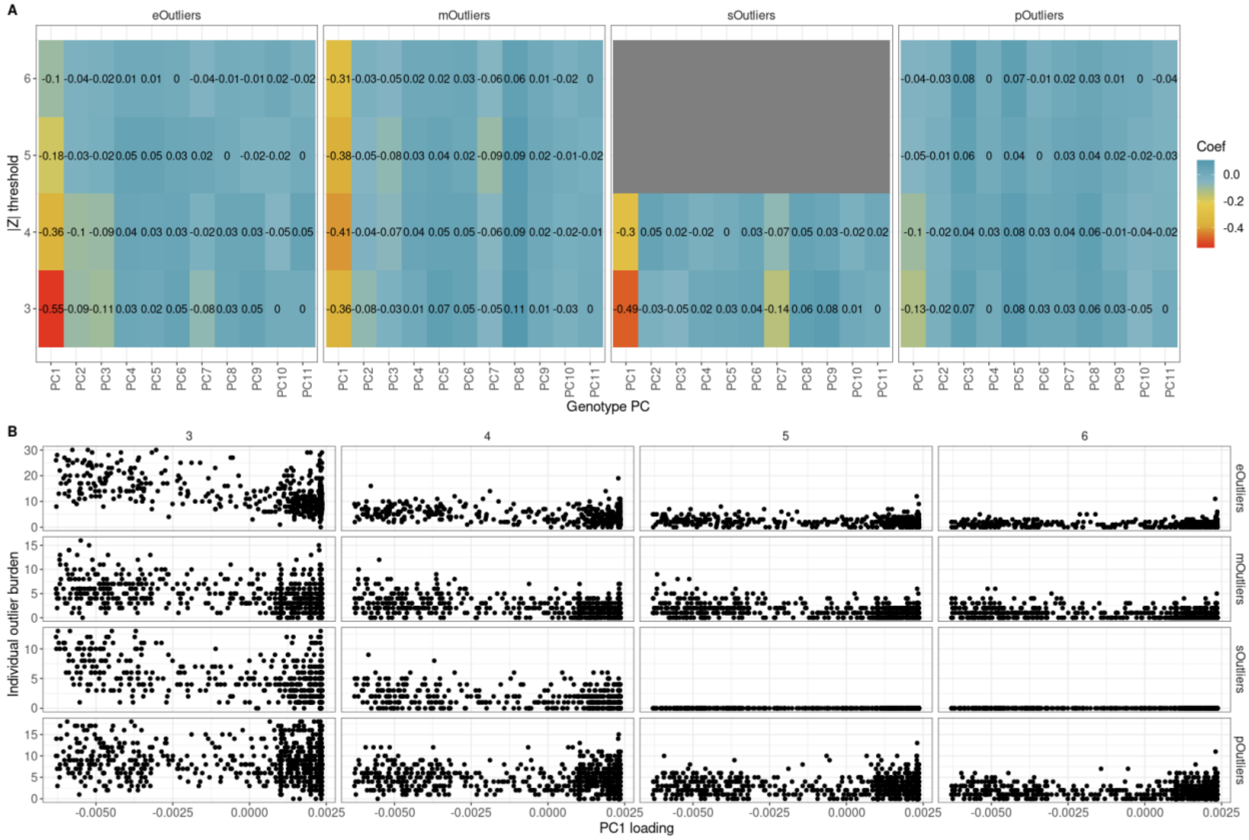


**Figure S8. Correlation of individual outlier burden and genotype PCs. (A)** The Pearson correlation of individual gene-level joint outlier burden across thresholds (y-axis) and individual genotype PC values (x-axis) for each gene-level outlier type. **(B)** Genotype PC1 values (x-axis) and individual outlier burden (x-axis) across thresholds (|Z| > 3, 4, 5, 6 in both exams) for each gene-level outlier type.

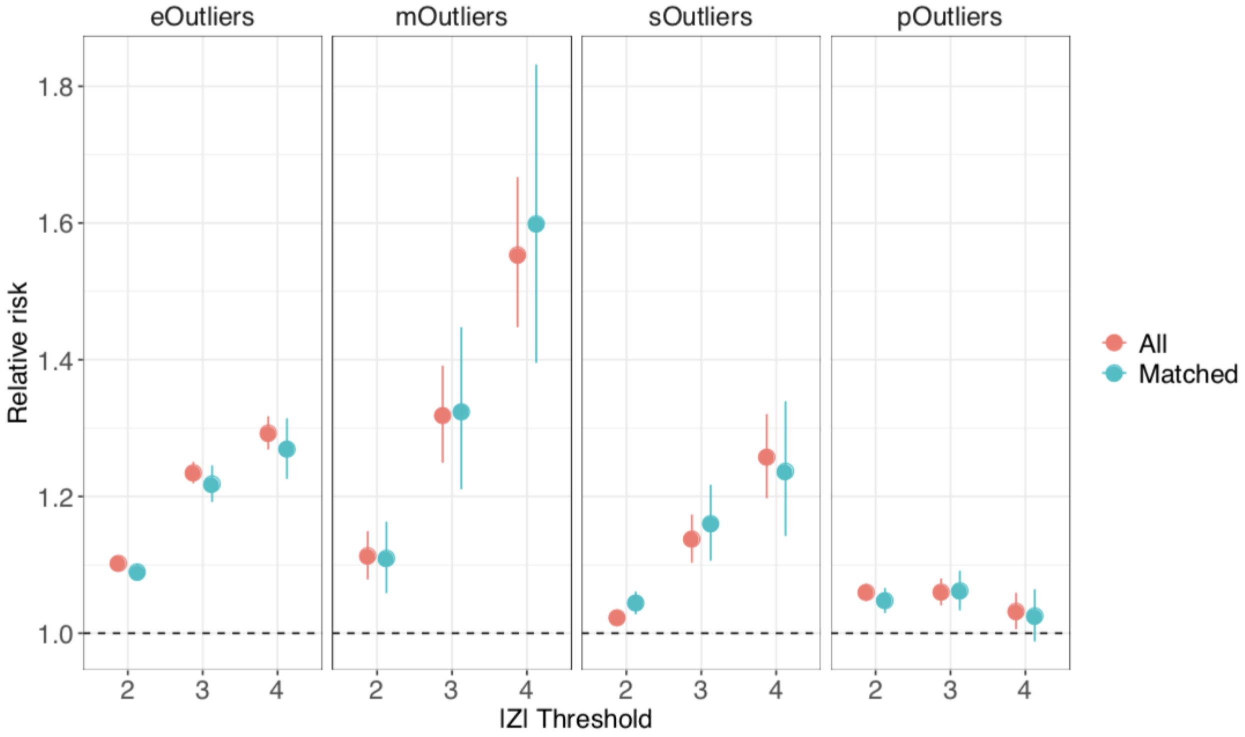


**Figure S9. Enrichment of rare variants nearby gene-level outliers relative to all controls or a subset matched by genotype PCs.** The relative risk of carrying a nearby rare variant (gene body +/- 10kb) across thresholds (x-axis) for gene-level joint outliers, considering as non-outliers all individuals with |Z| < 1 in both exams (pink) or selecting one non-outlier individual per outlier from all those with |Z| < 1 in both exams based on euclidean distance calculated across the top 11 genotype PC values (teal).

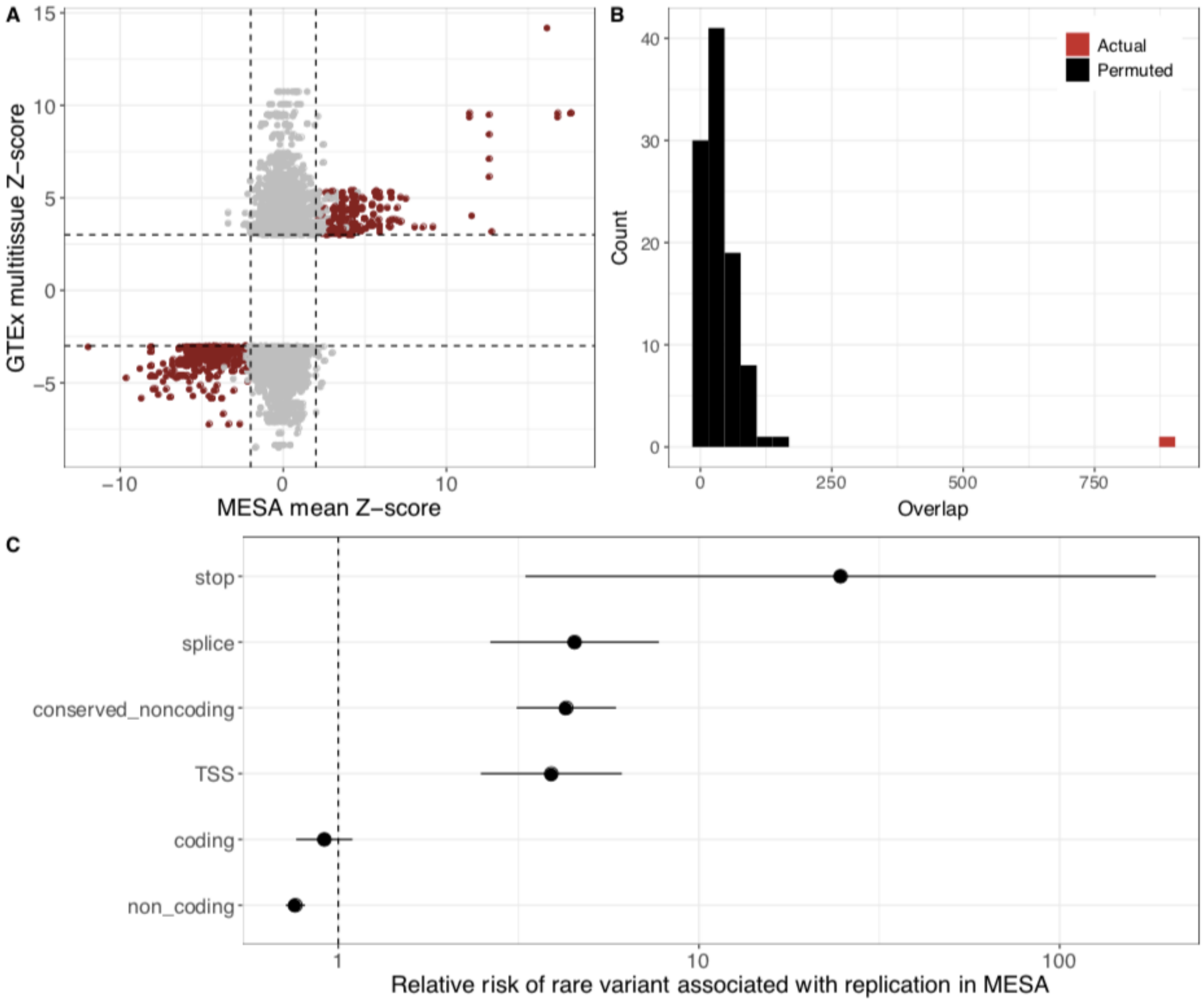


**Figure S10. Replication of GTEx multitissue-eOutlier associated rare variants in MESA. (A)** The mean expression Z-score across exams in MESA (x-axis) vs the median expression Z-score across tissues in GTEx (y-axis) for the set of variant-gene-individual instances where the same rare variant is carried by a GTEx and MESA individual and associated with multitissue outlier expression in GTEx. The dashed lines indicate the GTEx outlier threshold (|median Z| > 3) and the replication threshold in MESA (|Z| > 2 in both exams). Dark red indicates the instances that do replicate in both exams in MESA. **(B)** The number of instances in which the same rare variant is associated with outlier expression in GTEx and MESA (red), using a threshold of |Z| > 2 in both exams, vs the number observed after permuting expression Z-scores across all individuals in MESA, maintaining exam pairs. **(C)** The relative risk (x-axis) of a rare variant associated with outlier expression in GTEx replicating in MESA given that variant's annotation (y-axis). If a variant type was not associated with any replicating effect in MESA, it is not included here.

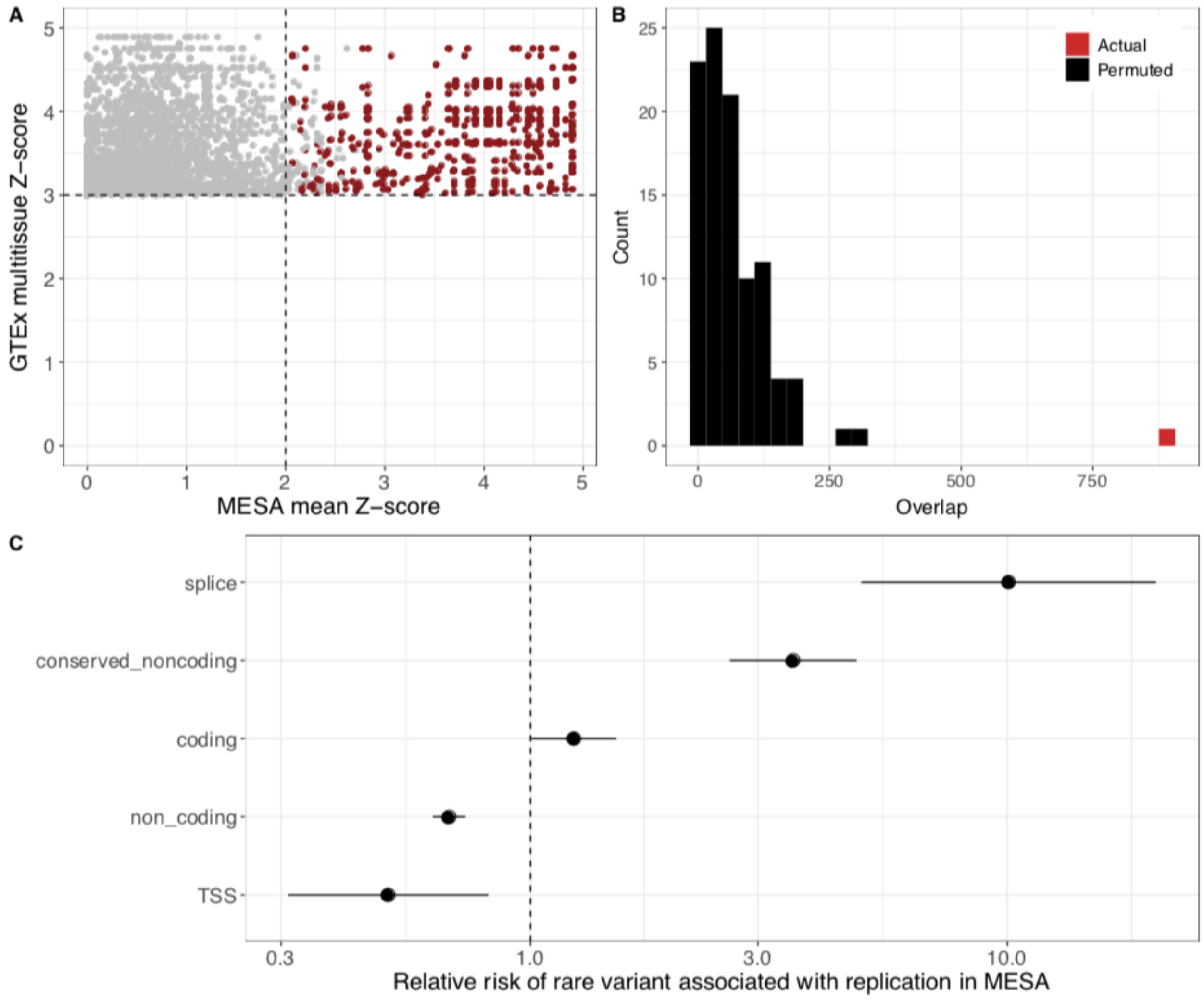


**Figure S11. Replication of GTEx multitissue-sOutlier associated rare variants in MESA. (A)** The mean splicing Z-score across exams in MESA (x-axis) vs the median splicing Z-score across tissues in GTEx (y-axis) for the set of variant-gene-individual instances where the same rare variant is carried by a GTEx and MESA individual and associated with multitissue outlier splicing in GTEx. The dashed lines indicate the GTEx outlier threshold (median Z > 3 or equivalently, median splicing p-value < 0.0027) and the replication threshold in MESA (Z > 2 in both exams). Dark red indicates the instances that do replicate in both exams in MESA. **(B)** The number of instances in which the same rare variant is associated with outlier splicing in GTEx and MESA (red), using a threshold of Z > 2 in both exams, vs the number observed after permuting splicing Z-scores across all individuals in MESA, maintaining exam pairs. **(C)** The relative risk (x-axis) of a rare variant associated with outlier splicing in GTEx replicating in MESA given that variant's annotation (y-axis). If a variant type was not associated with any replicating effect in MESA, it is not included here.

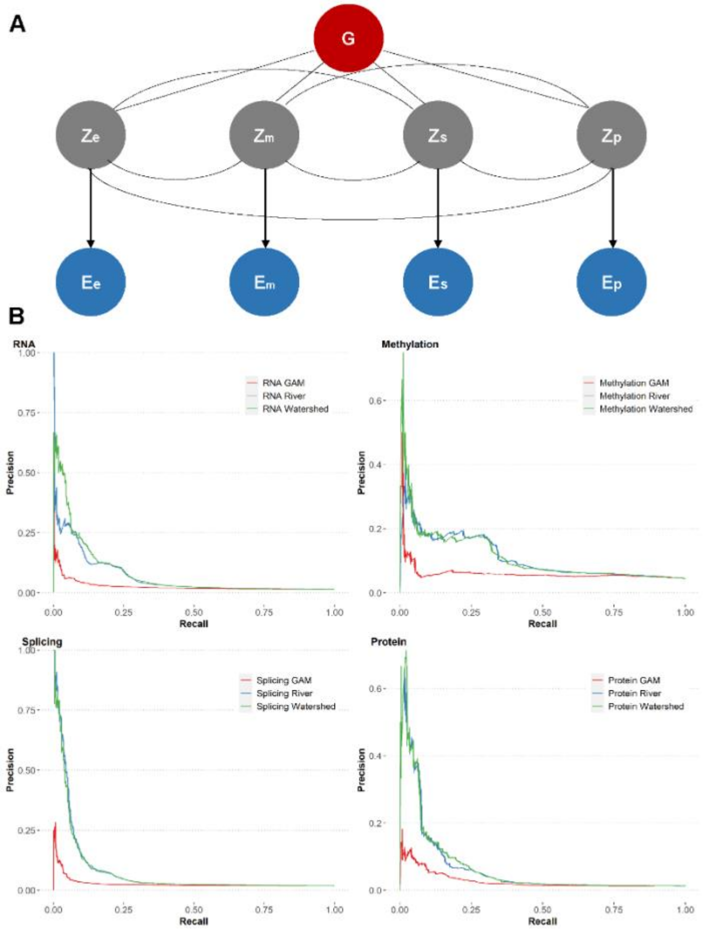


**Figure S12. Multi-omic Watershed model. (A)** Schematics of the Watershed hierarchical Bayesian model which is trained on (gene, individual) pairs consisting of genomic annotations aggregated across all rare variants (G), categorical variable E which represents observed outlier status of the gene in each signal (e – mRNA expression; m – methylation; s – splicing; and p – protein expression), and binary latent variables Z representing unobserved regulatory status on each signal. The Z layer is a fully connected layer which affords flexibility to model relationships across the regulatory cascade. **(B)** Precision-recall curves of Watershed models (green), River models (blue), and genomic annotation models (GAM, red) for each omic signal, evaluated against (gene, individual) pairs with the same set of rare variants nearby.

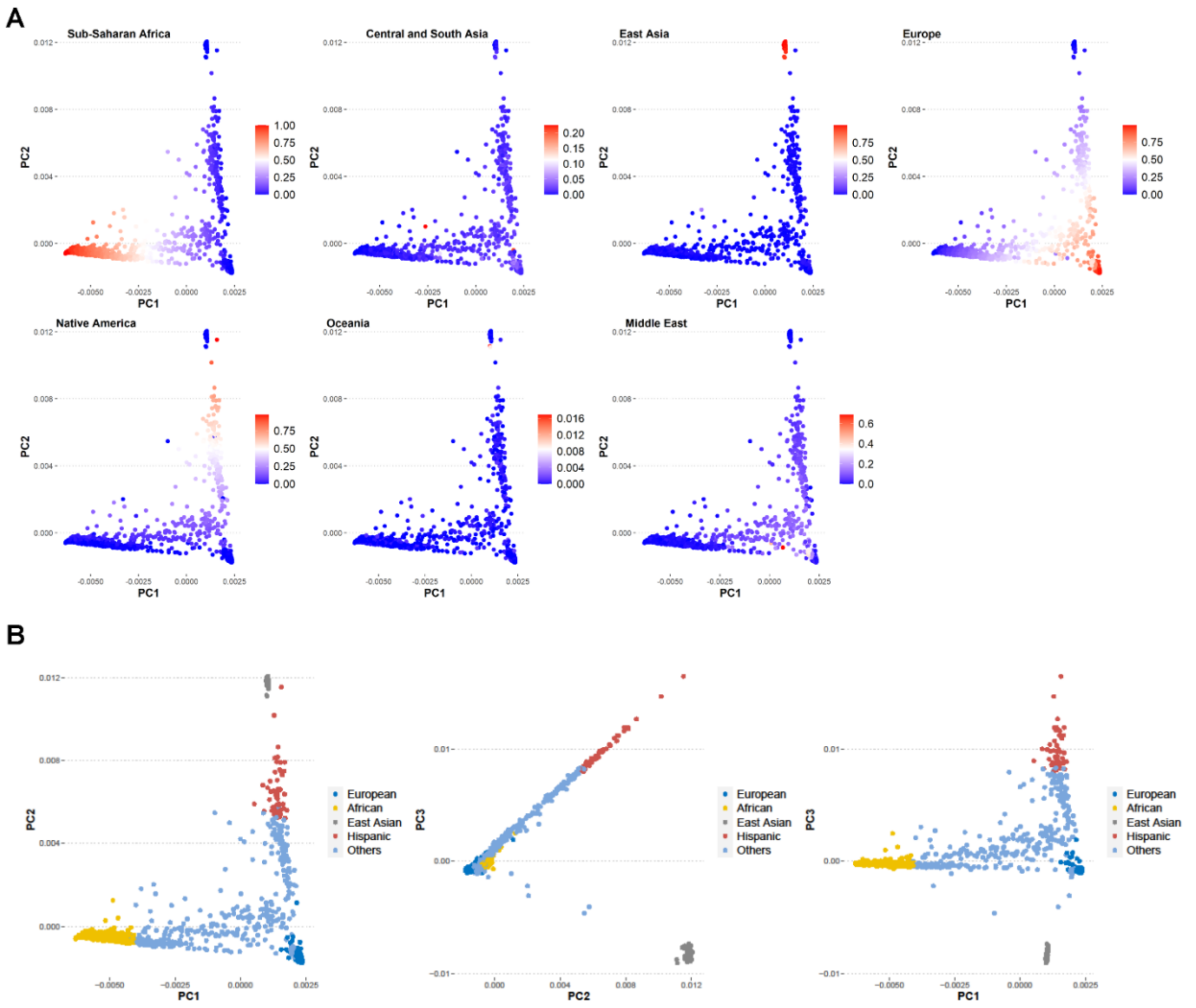


**Figure S13. Population diversity in MESA. (A)** Genotype principal component (PC) plots for N = 1319 individuals in MESA with multi-omic measurements, colored by inferred ancestry using RFMix (Maples et.al. 2013) based on the Human Genome Diversity Panel (HGDP) with seven super populations. **(B)** Genotype PC plots for the same set of individuals colored by population grouping obtained by setting thresholds on inferred ancestry in (A). The resulting populations have N = 426 Europeans, N = 270 Africans, N = 107 East Asians, and N = 54 Hispanic individuals.

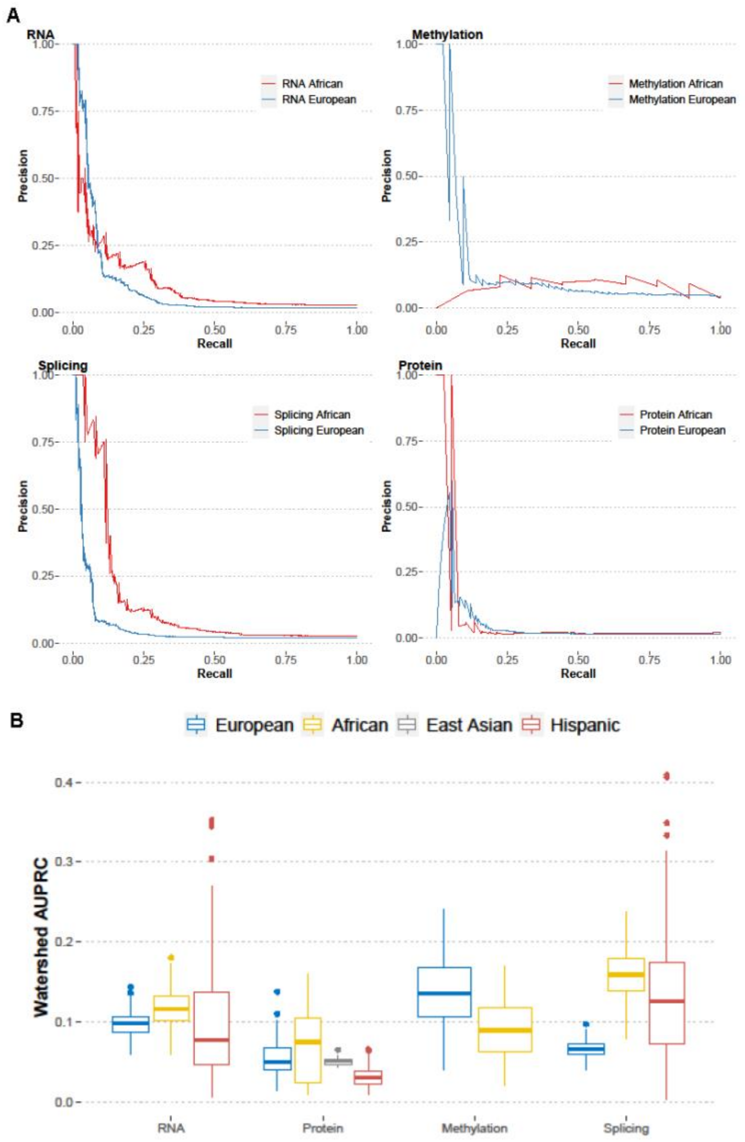


**Figure S14**. **Cross-population Watershed performance**. **(A)** Precision-recall curves for Watershed models trained on European individuals and evaluated on either European N2 pairs (blue) or African N2 pairs (red) across four omic signals. **(B)** Summary of 100 bootstrapped area under the precision-recall curves (AUPRC) for Watershed models trained on European individuals and evaluated on other populations in MESA across our omic signals. Shown are median and interquartile ranges. Only populations and signals with at least ten pairs of N2 individuals for evaluation are included.

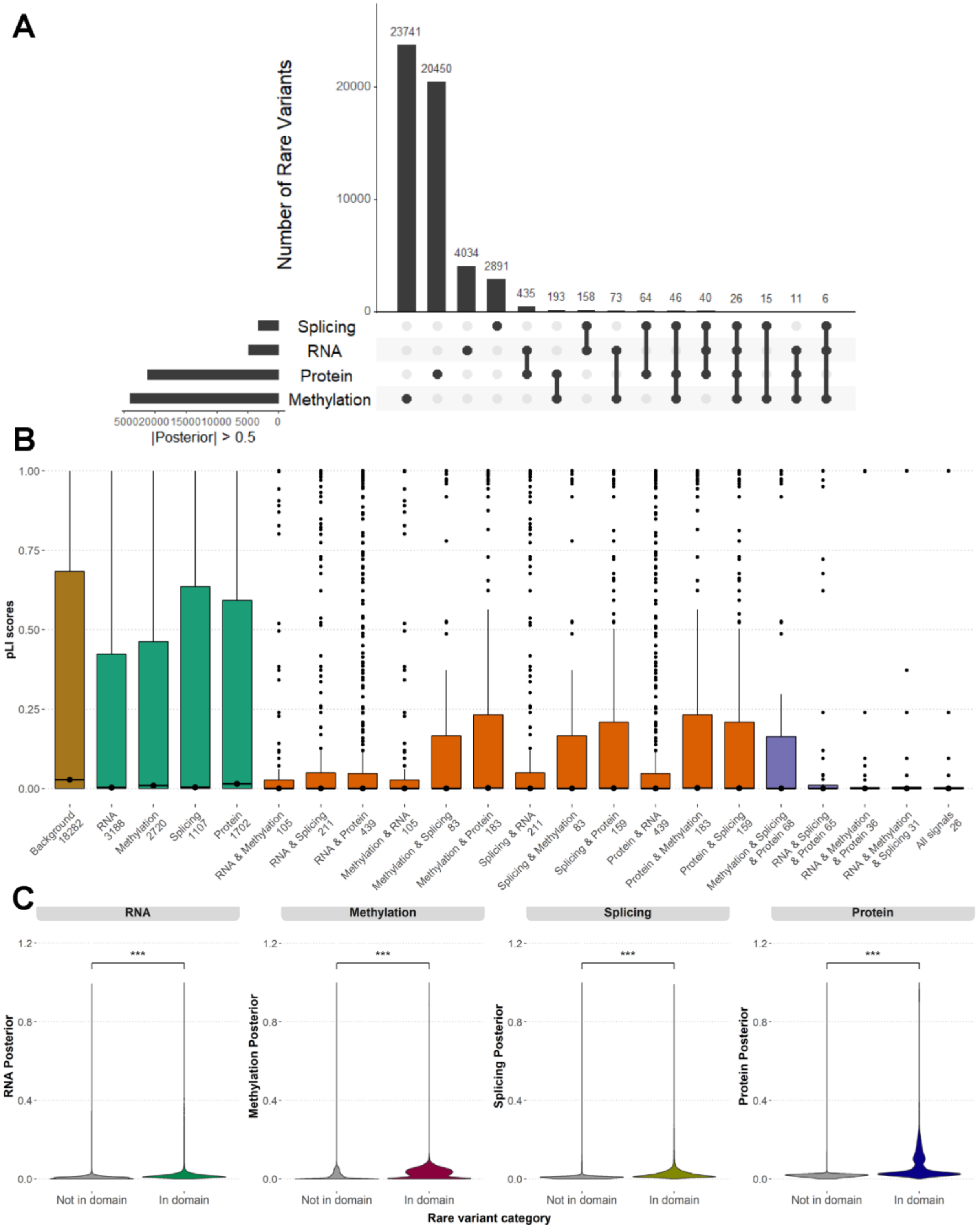


**Figure S15. Cross-modality comparison of Watershed posteriors**. (A) Upset plot of all rare variants passing 0.5 posterior threshold in each omic signal. (B) Distribution of probability of loss of function intolerance (pLI) for genes with rare variants passing 0.5 posterior threshold in each signal individually (green), and combinations of two (orange), three (purple), and all four (red) signals. Background is represented by all genes with any omic measurement in MESA. Number of rare variants in each group is shown in the x-axis labels. Median and interquartile range is shown for each distribution. (C) Violin plots of Watershed posteriors in each signal for rare variants identified in protein domains, compared with those not in domains. *** $p < 0.001$, one-sided Wilcoxon rank-sum test on absolute value of posteriors between the two variant category.

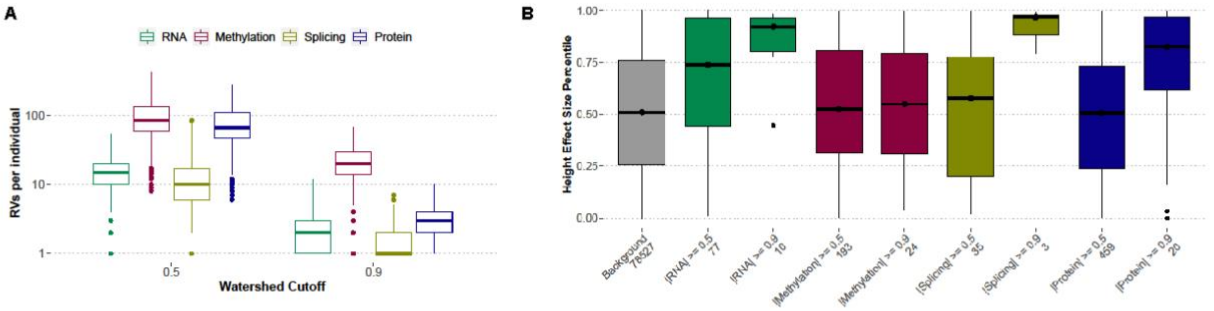


**Figure S16. Assessment of Watershed performance for genes directly measured in each omic signal.** (A) Number of rare variants per individual as prioritized by each omic signal at two levels of Watershed posterior cutoff 0.5 and 0.9. Individuals with a significantly large number of outlier expressions ("global outliers") are excluded. Similar to Figure 5A, except only genes which are directly measured in each omic signal are included. (B) Distribution of percentile normalized effect size for height (median and interquartile range) of all rare variants (background, gray), and those variants prioritized by Watershed in each signal. Only those rare variants mapped to genes with evidence of causing abnormal body height which are directly measured in each omic signal are included.

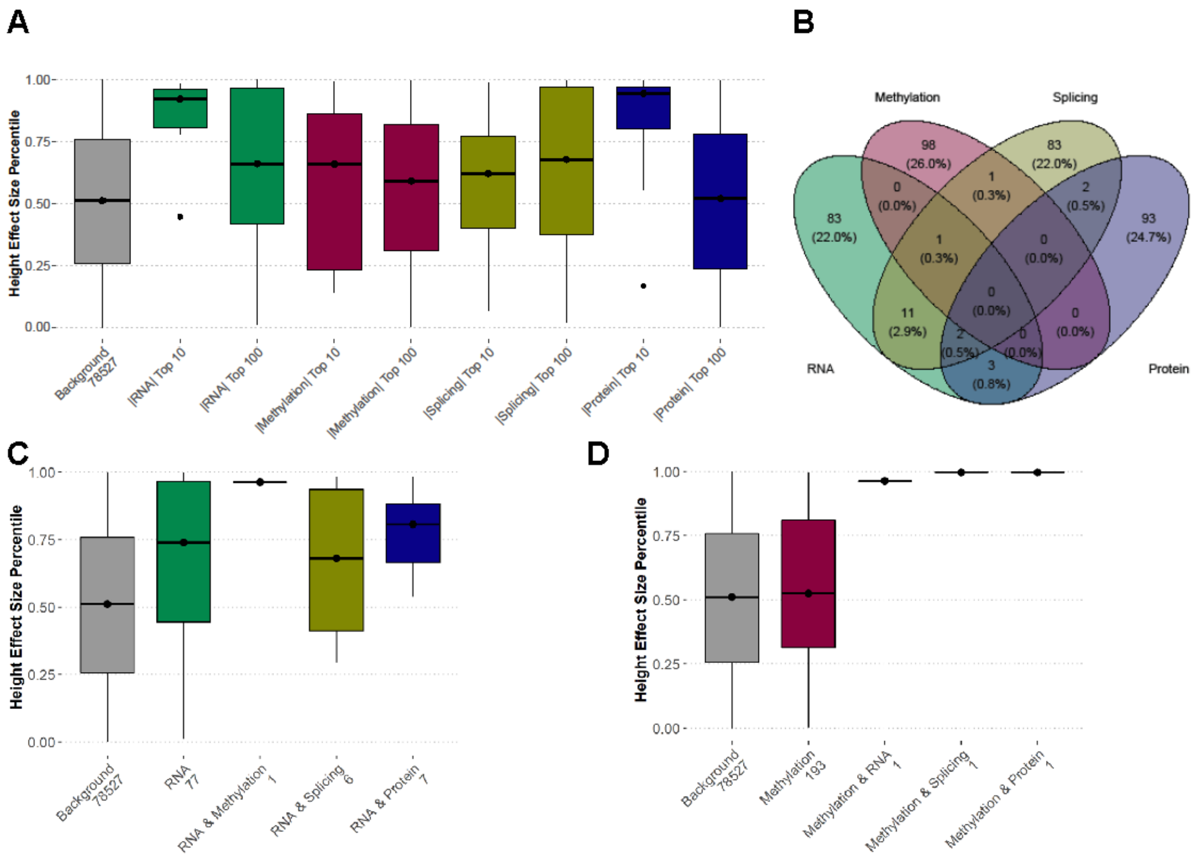


**Figure S17. Evaluation of the effect size for height for rare variants prioritized by the multi-omic Watershed model.** (A) Distribution of percentile normalized effect size for height (median and interquartile range) of all rare variants (background, gray), and those rare variants prioritized by multi-omic Watershed in each signal with top 10 or top 100 highest posteriors. (B) Venn diagram of top 100 rare variants with highest posteriors across signals. (C – D) Distribution of percentile normalized effect size for height (median and interquartile range) for rare variants prioritized by a single signal at a posterior threshold of 0.5 (C – RNA; D – methylation) and combined with another signal.

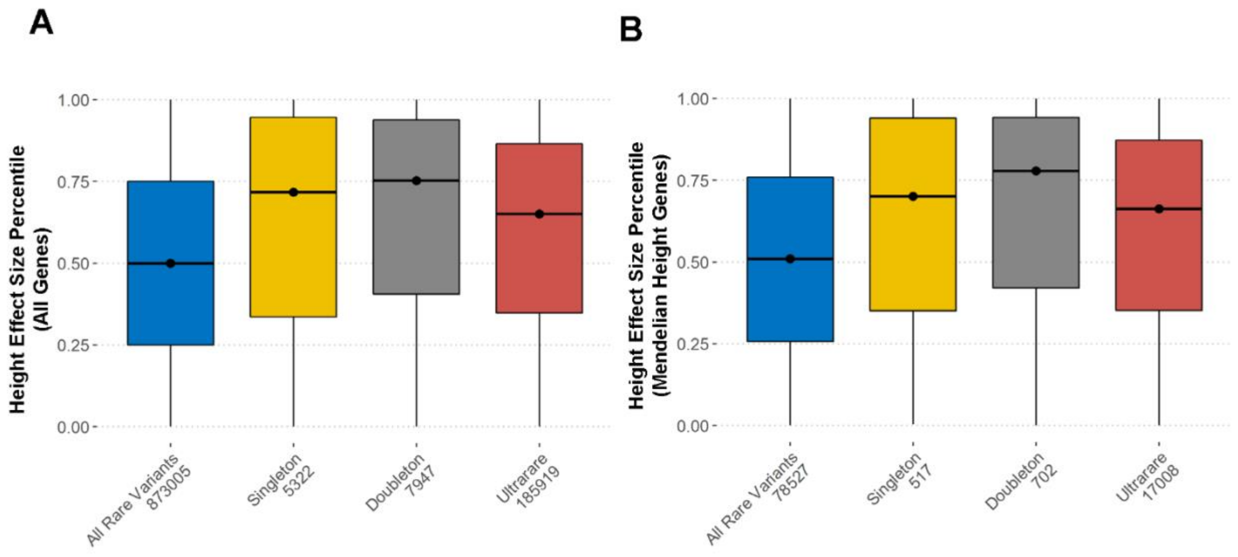


**Figure S18. Evaluation of the effect size for height by minor allele frequency (MAF)**. Distribution of percentile normalized effect size for height (median and interquartile range) for all rare variants (blue), singletons (yellow), doubletons (grey), and ultrarare variants (MAF < 0.1%) across all genes **(A)** or N = 1,314 genes with evidence causing abnormal body height (Mendelian height genes, **B**).

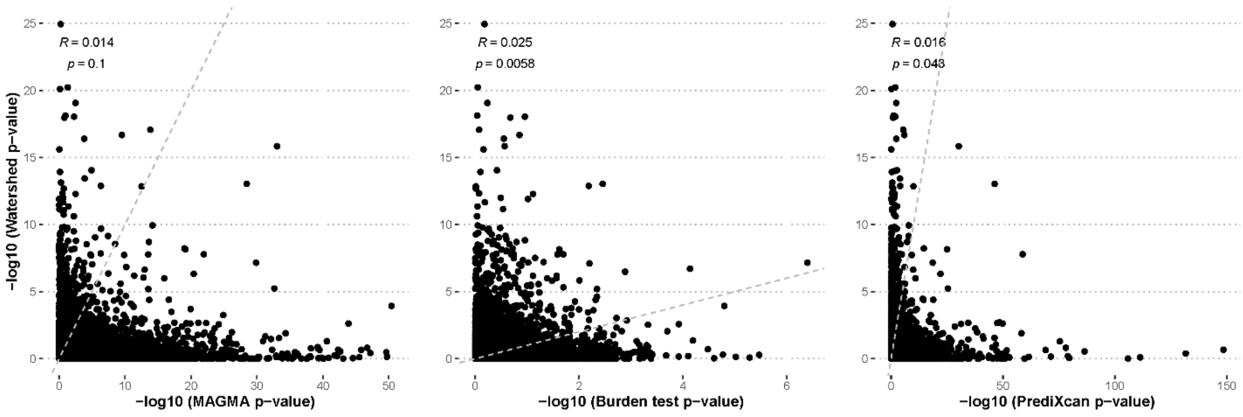


**Figure S19. Comparison of collapsing analysis based on Watershed posteriors with other gene prioritization methods.** Scatter plots of -log10 p-values of gene tests using Watershed posteriors as weights (y-axis) against those obtained from MAGMA **(A)**, burden test **(B)**, and PrediXcan **(C)** on height. Dotted lines represent the diagonal.

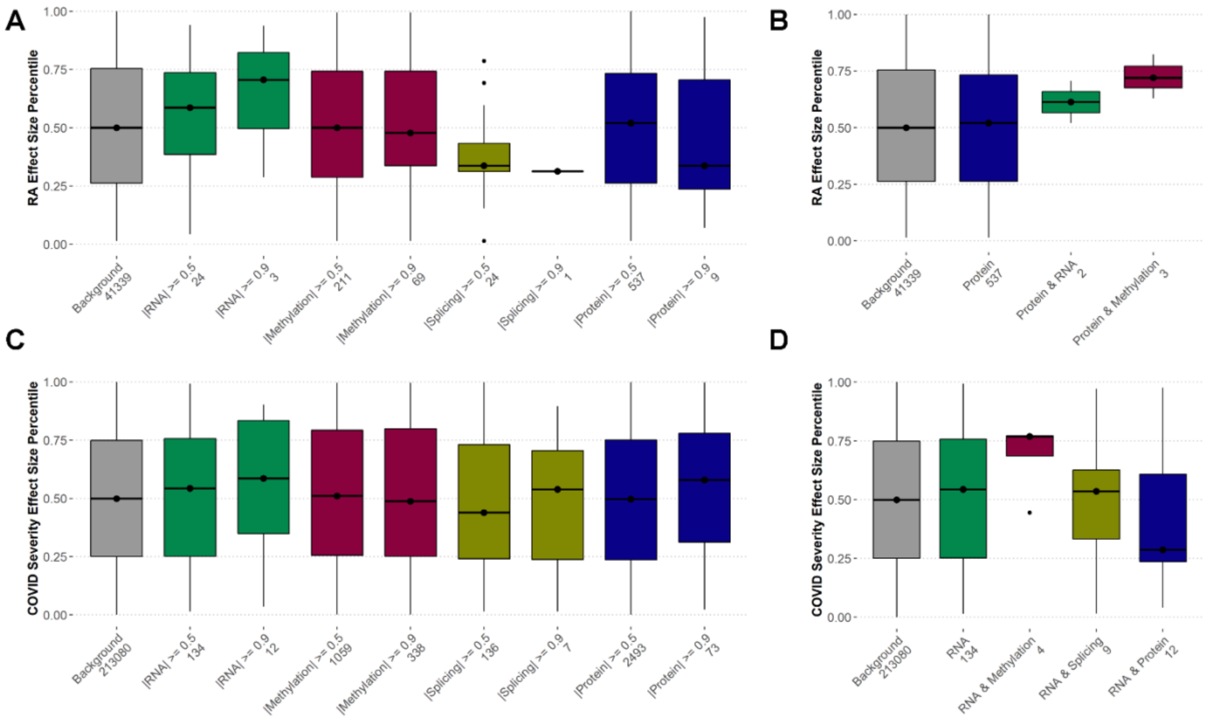


**Figure S20. Evaluation of Watershed prioritized rare variants in two immune diseases. (A)** Distribution of percentile normalized effect size for rheumatoid arthritis (RA, median and interquartile range) for all rare variants (background, grey), and those rare variants prioritized by multi-omic Watershed in each signal at two posterior threshold values. Only rare variants mapped to genes with evidence of association with RA as reported by Open Targets are shown (N = 4,769 genes). **(B)** RA effect size distribution as in **(A)** but with rare variants prioritized by protein (blue) or protein combined with another signal at a posterior threshold of 0.5. **(C)** Distribution of percentile normalized effect size for COVID-19 severity for all rare variants, and those rare variants prioritized by multi-omic Watershed in each signal at two posterior threshold values. Only rare variants mapped to genes with evidence of association with COVID-19 as reported by Open Targets are shown (N = 2,232 genes). **(D)** COVID-19 severity effect size distribution as in **(C)** but with rare variants prioritized by RNA (green) or RNA combined with another signal at a posterior threshold of 0.5.

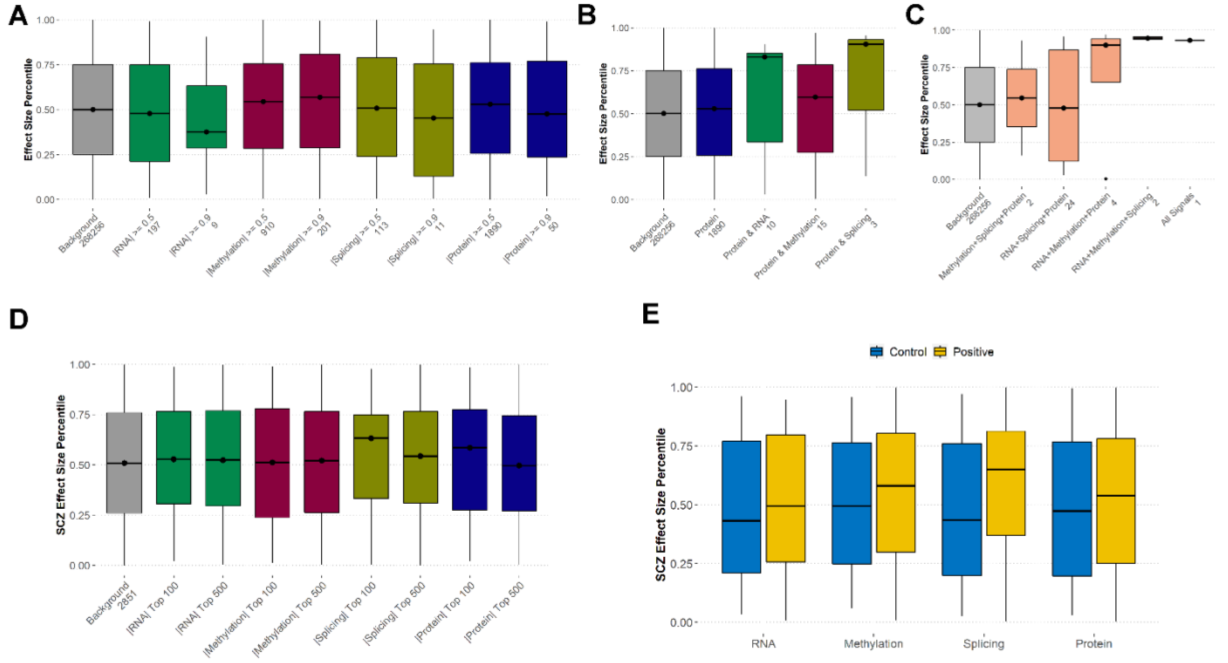


**Figure S21. Evaluation of Watershed prioritized rare variants in two neurological diseases. (A)** Distribution of percentile normalized effect size for Alzheimer's Disease (AD, median and interquartile range) for all rare variants (background, grey), and those rare variants prioritized by multi-omic Watershed in each signal at two posterior threshold values. Only rare variants mapped to genes with evidence of association with AD as reported by Open Targets are shown (N = 7,103 genes). **(B)** AD effect size distribution as in **(A)** but with rare variants prioritized by protein (blue) or protein combined with another signal at a posterior threshold of 0.5. **(C)** AD effect size distribution as in **(A)** but with rare variants prioritized by three or four omic signals. **(D)** Distribution of percentile normalized effect size for schizophrenia (SCZ, median and interquartile range) of all rare variants (background, gray), and those rare variants prioritized by multi-omic Watershed in each signal with top 100 or top 500 highest posteriors. Only rare variants mapped to genes with evidence of association with AD as reported by Open Targets are shown (N = 5,463 genes). **(E)** Distribution of SCZ effect size of rare variants mapped to MAGMA prioritized genes ("Positive", MAGMA z > 2, N = 5,378 genes, yellow) and control genes ("Control", MAGMA z < 0, N = 4,092 genes, blue) at a posterior threshold of 0.2 in each omic signal.
